# Merkel cell polyomavirus infection and persistence modelled in skin organoids

**DOI:** 10.1101/2025.02.11.637697

**Authors:** Silvia Albertini, Manja Czech-Sioli, Thomas Günther, Sanamjeet Virdi, Patrick Blümke, Lisann Röpke, Veronika Brinschwitz, Maura Dandri, Lena Allweiss, Rudolf Reimer, Carola Schneider, Arne Hansen, Susanne Krasemann, Emanuel Wyler, Markus Landthaler, Adam Grundhoff, Nicole Fischer

## Abstract

Merkel cell polyomavirus (MCPyV) causes most Merkel cell carcinomas (MCC). The virus is one of the few known human tumor viruses, and due to its direct role in this skin cancer development, it is a bona fide model for viral tumorigenesis and tumorigenesis in general. Chronic viruses in humans such as MCPyV are highly adapted to their host and current models to study infection, persistence and pathogenesis are highly limited. We here use an induced pluripotent stem cell (iPSC)-derived hair-bearing skin organoid (SkO) system to demonstrate efficient viral infection, progression and spread of MCPyV. Using bulk-, single cell - and spatial-transcriptomics, combined with immunostaining and nucleic acid hybridization technologies, we show that MCPyV ensures persistence due to a quasi-latency state within the majority of dermal fibroblasts carry the viral genome. Further, we identify the cell type of productive infection with papillary fibroblasts and dermal sheath fibroblasts supporting viral replication and progeny production. Our high-resolution methods demonstrate that the virus in these cells evades the innate immune response, as evidenced by the efficacy of interferon-beta treatment or ruxolitinib, a JAK/STAT inhibitor, in suppressing or stimulating viral replication. We show that iPSC-derived SkOs are able to support infection and long-term persistence of the virus under conditions very similar to those found in humans. Thus, this infection model provides a robust platform for understanding and characterizing the interaction of this virus with the immune system in infection, testing treatment strategies to control reactivation and map processes involved in tumor development.

## Introduction

Merkel cell polyomavirus (MCPyV) has been identified as the causative agent for the majority of Merkel cell carcinomas (MCC), accounting for up to 80% of MCC cases ^1–5^. It is one of seven human tumor viruses that have been identified to date ^2^. MCPyV was first detected in 2009 through a comprehensive analysis of the transcriptome in primary cases of MCC ^3^. This particular tumor has been observed to predominantly affect elderly or immunocompromised patients ^3^. In contrast to other human tumor viruses, which are known to cause tumors through constant reinfection of cells and thus long-term chronic inflammation (for example, hepatitis B virus, HBV, and hepatitis C virus, HCV), or by oncogenic transformation out of viral latency in conjunction with co-factors such as Epstein-Barr virus, EBV, and Kaposi-Sarcoma virus, KSHV, MCPyV is unique in its direct tumor etiology ^2–7^. MCCs that are positive for the tumor are monoclonal, with each tumor cell carrying the viral genome integrated into the host genome. The tumor cell viability is dependent on viral oncogene, small T-Antigen (sT) and large T-Antigen (LT), carrying tumor-specific mutations, expression ^4,5^.

MCC is a poorly differentiated neuroendocrine carcinoma that arises in the dermis, progresses rapidly, and is divided into two molecular subclasses ^1^. These two classes, MCPyV-positive MCCs and UV-associated virus-negative MCCs, differ significantly in the mutational burden of the host genome. Virus-negative MCCs have a 100-1000-fold higher mutational load compared to virus-positive tumors, with many oncogene-activating events and transcriptional differences ^6–8^. MCPyV is an opportunistic pathogen that is asymptomatic in the immunocompetent host, highly controlled by the immune system, and reactivates and replicates in immunosuppressed individuals, leading to severe disease through mutation in the viral early region and integration of the viral genome. The virus is highly prevalent in the general population, with more than 80% of adults positive for MCPyV-specific antibodies, however MCC disease is a rare with an incidence of 0.7-1.6/100,000 ^1^.

The virus is a non-enveloped, double-stranded, circular DNA virus which, due to its small size of 5kbp, encodes only a few proteins. The early gene region encodes LT, sT, 57kT and ALTO, an alternative LT open reading frame, and a viral microRNA, miR-M1. The late region encodes two nucleocapsid-forming proteins VP1 and 2 ^2,9^. The virus depends on the host’s DNA replication machinery for its reproduction. The infection systems developed for the virus thus far, such as through the transfection of a recombinant virus genomes, have yielded only a limited number of virus-positive cells, amounting to less than 5% positive cells in monolayer cultures. These virus positive cells do not mount the entire viral life cycle, but rather only early gene expression and virus DNA replication ^10–12^. However, MCPyV particles can be produced in low amounts in helper cells that overexpress sT and LT ^11,13^. These MCPyV particles have been utilized to infect various cell types, including fibroblasts ^11,13,14^. Similarly to re-ligated transfected genomes, the infection rate is low, with only a few cells positive for LT and VP1. It is noteworthy that when fibroblasts are freshly isolated from biopsies and stimulated to proliferate with various growth factors, the infection rate can be increased to 20%, although no virus progeny, released virus, is produced ^11^. Although PyVs demonstrate considerable similarities to papillomaviruses, e.g., exhibiting high cell type restriction, there are significant differences between these viruses. These differences are reflected, in part, by the fact that air-liquid interface (ALI) cultures, which consist of fibroblasts and layers of differentiated keratinocytes found in the epidermis, cannot model MCPyV infection. To understand the processes of infection, chronic persistence, immune control of the virus and the first steps of tumorigenesis, an infection model is essential that can mimic the course of infection in humans. Surrogate models, such as mouse models that reflect infection and tumor initiation do not exist due to the restricted host tropism ^15–17^.

The development of organoids from tissue derived-derived stem cells or induced pluripotent stem cells ^15–17^ is a rapidly advancing improvement in the representation of physical and physiological processes in tissues during development and especially in the perturbation of the normal state, such as by infection. This has been demonstrated in some studies in the past, particularly in two studies of emerging infections, SARS-CoV-2 and Mpox ^18,19^. Both are acute infectious agents that have very short infection cycles and do not cause long-term persistent infections. Fundamental questions about these viruses, such as the infection cycle, can be addressed in simple cell culture systems. In our study, we show that human iPSC-4 derived SkO models, which have been recently developed ^20,21^, represent bona fide models to depict persistent infections; we here show that we can model the life cycle of the human tumor virus MCPyV over 120 days, map its interaction with innate immunity, and identify the different cell types that contribute to infection. We show that complex tissue models require the use of advanced imaging and single cell analysis, both temporally and spatially, to map and understand the course of infection.

The development of complex tissue models derived from human iPS cells, including skin organoids, represents a promising approach to elucidating the infection cycle of challenging human persistent infection-causing viruses and comprehending their life cycle. This model holds significant potential in facilitating the development of intervention strategies for both chronic infection and pathogenesis.

## Results

### MCPyV efficiently infects human iPSC-derived Skin Organoids (SkOs)

Currently available *in vitro* infection systems for MCPyV are limited to semi-permissive infection systems that transfect re-circularized viral genomes into cell lines and result in replication of viral DNA and viral gene expression in few cells for short periods of 2-6 days only ^10,12,13^. Interestingly, using this model, it was demonstrated that the MCPyV genomes can persist long-term in culture independent of re-infection ^22^. In addition, an i*n vitro* infection system was described involving infection of human dermal fibroblasts grown in monolayers ^11,13,14^. The system allows only few cells to be infected and followed for short time periods. These models do not support new virus particle release or spread of infection, which hinders the use of these systems for studying the life cycle, cell tropism and initial pathogenesis mechanisms. We have generated hair-bearing skin organoids (SkOs) from human induced pluripotent stem cell lines (hiPSC line UKEi001) according to previously published protocols and assessed the correct differentiation over time applying whole-mount immunostaining (WMI) as described recently ^20,21^, (Supplementary Figure S1).

Concurrent with previous reports^18,20,23,24^, we observe iPS cells forming a translucent cyst starting from day 8 of differentiation (dod) and a polarized conformation around 30-45dod, with a dense tail containing the ganglia of the neurons (TUJI^+^) and a head characterized by stratified epithelium (KRT15^+^, KRT17^+^, and ECAD^+^) surrounded by dermal fibroblasts (PDGFRα^+^). We further observe Merkel cells, distributed in the basal layer of the epithelium (KRT20^+^), at 30dod, increasing in frequency overtime and predominantly located in the bulge area of the hairs 120dod. Hair placodes (SOX2^+^ and LHX2^+^) initiate around 65dod. Organoids were considered fully matured when displaying innervated hair follicles (>120dod, Figure S1b, c). We infected fully differentiated SkOs by adding MCPyV (10E8 293-MCPyV genome equivalents), produced in helper cells, to the SkO culture supernatant. We collected SkOs at various time points, extending up to 30 days post-infection (dpi). We performed WMI for MCPyV capsid protein (VP1), in conjunction with KRT17 (a marker for periderm, epidermis, and outer root sheath of the hair follicle), thereby revealing few VP1-positive cells in the outer layer of the organoid, the site where predominantly dermal fibroblasts are located. The frequency of VP1 positive cells slightly increased over time (Figures S2a, b). In fully differentiated SkOs, which due to their inset-out morphology have already accumulated a large number of detached squamous epithelial cells inside the organoid, the maximum observation time is limited. In order to observe the progression of the infection over an extended time period, we infected SkOs 30-45dod. Longitudinal observation of the infection with different organoids sacrificed at specific time points after infection demonstrates that the number of VP1-positive cells in the outer layer of the dermis is initially low (Figures S2c, d and S3b), however, increases significantly over time (Figure 1b and Supplementary Figure S2d and S3b), reaching more than 6000 VP1-positive cells/mm³ at 120dpi. MCPyV VP1 staining was observed exclusively in fibroblasts within the dermis (Figures 1a and Supplementary Figures S2d and S3b), while VP-1 staining was not detected in epidermal cells, identified using KRT17 as a marker. The results of the early and late time points demonstrate a consistent pattern, with an increase in the number of positive cells over time, starting from the outer layer and progressively spreading into the more central layers of the dermis (Supplementary Figures S2 and S3). The observed differences in the total number of VP1-positive cells at all time points starting from 15 days post-infection (dpi) to 90dpi in SkOs 30dod or 150dod (see Supplementary Figure S3b) can be attributed to the increased diffusion capacity of MCPyV in the early infected organoid 30dod. The dermis of SkOs at early stage of differentiation is distinguished by a reduced extracellular matrix compared to late-differentiated SkOs. We confirmed this by DNAScope analysis using MCPyV-specific probes demonstrating that SkOs infected at 30 vs. 135dod and followed for 15 days clearly demonstrate an increased DNAScope signal in SkOs at earlier differentiation states, such as 30dod (Supplementary Figure S4). During the course of the infection, the organoid culture media was collected and examined for MCPyV viral copy numbers using quantitative PCR (Figure 1c). After an initial decrease in viral copy number, which represents the dilution of the virus by the medium change, we observe an increase in viral genome copies starting from 27dpi, which continues to increase over the further course. The culture medium of the SkOs 27dpi and later was pooled and concentrated by ultracentrifugation (SkO-MCPyV). The samples were then examined morphologically and in re-infection experiments in comparison to the 293-MCPyV virus as a control. The application of negative staining electron microscopy to the concentrated virus confirmed the presence of both full and empty virus particles, with an approximate size of 40–60 nm (Supplementary Figure S5a) with no significant differences between the two viral populations. Furthermore, re-infection experiments on both monolayer fibroblasts (nHDFs) and SkOs infected with either SkO-MCPyV or 293-MCPyV confirmed the infectivity of both viruses (Supplementary Figures S5b and S5c), which did not significantly differ in infection efficiency or cell types infected.

**Fig. 1.**
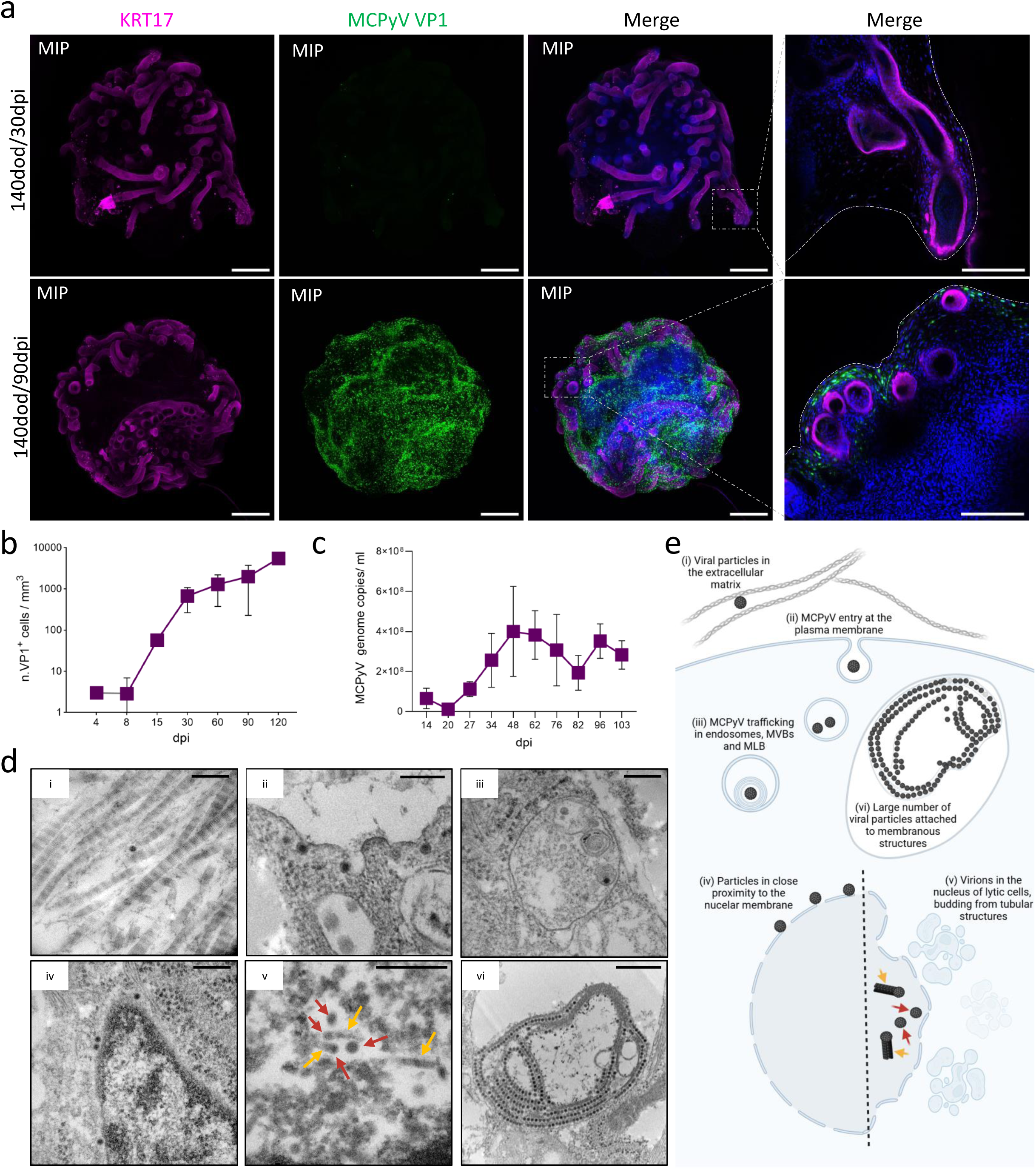
MCPyV establishes a productive and spreading infection in human iPSC-derived skin organoids. **a,** Whole-mount immunostaining of 140 days of differentiation (dod) SkOs infected for 30 days post-infection (dpi) or 140dod/90dpi with 1.5×10E08 genome equivalents of MCPyV per SkO. KRT17 staining for epidermis and outer root sheath of the hairs (magenta), MCPyV VP1 staining for the virus capsid protein (green), and Hoechst counterstaining for nuclei (blue). Where indicated, the images show the maximum intensity projection (MIP) of multiple z-stack acquisitions. Elsewhere a single z-stack is shown. Dashed boxes indicate the magnified regions on the right. The dashed line in the single z-stack magnification indicates the SkO outer margin. Scale bar: 500 μm for the main images, 200 μm for the magnifications. **b,** Quantification of VP1 positive cells per mm^3^ of SkOs infected at day 30 of differentiation and fixed at the indicated days post infection (dpi, x-axis). Values are presented as mean of biological replicates ± s.e.m (n = 2). **c,** Quantitative real-time PCR showing the copies of viral DNA in the SkO supernatant over time. Values are presented as mean of biological replicates ± s.e.m (n = 4). **d,** Representative transmission electron microscopy (TEM) images showing 40-60 nm electron-dense particles in a 150-day-old organoid infected with MCPyV for 100 days. Scale bar: 200 nm in i-v, 500 nm in vi. **e,** schematic representation of the TEM results shown in panel d (created with BioRender.com).

In addition to the SKO culture medium, we examined the SkO tissue infected for 100 days for virus particles and structures indicative of infection using transmission electron microscopy (TEM). We find electron dense particles representative of 50–60 nm MCPyV in the dermis of SkOs (Figure 1d and Supplementary Figure S6). The presence of MCPyV particles was observed within the extracellular matrix, in close proximity to collagen fibers (Figure 1d, panel i), within invaginations of the plasma membrane (Figure 1d, panel ii), and within endosomal compartments of multivesicular and multilamellar bodies (Figure 1d, panel iii) both representing entry, incoming particles ^25^. We find viral particles at the nuclear lamina, suggesting that they leave the endoplasmic reticulum (ER) and are transported through the cytosol before entering the nucleus (Figure 1d, panel iv) ^26–28^ and in tubular structures similar to those of murine polyomavirus (Figure 1d, panel v and Supplementary Figure S6d), which could be related to virus formation in the nucleus^29^. Of particular interest are the large clusters of virus particles observed within endosomal compartments, which have an approximate diameter of 2 µm (Figure 1d, panel vi and Supplementary Figure S6e). These structures could potentially be remnants of cell nuclei from neighboring cytopathic cells, as analogous structures have been observed in the extracellular matrix of SkOs (Supplementary Figure S6e). Overall, our findings demonstrate that MCPyV establishes a productive and spreading infection in SkOs in dermal fibroblasts.

### Single-cell and spatial transcriptome analyses reveal different fibroblast subpopulations in the organoids

Our observation that a substantial proportion of fibroblasts within the dermis carry the viral genome, yet only a limited number of these cells also exhibit detectable protein expression of the viral early protein LT antigen as shown by DNAScope and immunohistochemistry (Supplementary Figure S4), prompted us to undertake a more comprehensive characterization of the fibroblast populations within the dermis. We conducted a multi-omics analysis of MCPyV-infected SkOs, a schematic overview of the organoids and time points utilized for the various sequencing techniques is depicted in Supplementary Figure S7. The lack of publicly available sequencing data on fully differentiated SkOs and their representative cell types, or, given that SkOs represent fetal skin in the second trimester at the differentiation stage ^20^, on samples of human embryonic skin, prompted us to combine data from single-cell RNA-sequencing (scRNA-seq) with data received by spatial transcriptomics to identify the cell types represented in SkOs. UMAP dimensionality reduction of scRNA-seq revealed several distinct clusters that were manually assigned to each skin population based on the expression of known markers ^20,30–32^ (Figure 2a-b and Supplementary Figure S8 and S9). The vast majority of SkO cell types in scRNA-seq were attributable to fibroblast populations (enriched for PDGFRα, MEST and COL5A2). This is partly due to the use of fully differentiated SkOs in the single-cell analysis, which are very rigid at these late stages of differentiation. Compared to the tightly associated keratinocytes and hair follicle cell types, fibroblast cell layers are relatively easy to dissociate due to their loose association and location in the outermost layer of the SkOs. However, a comparison of the size of the fibroblast populations in scRNA-seq and spatial transcriptomics reveals that fibroblasts are the predominant cell population in SkOs. Specifically, scRNA-seq analyses indicated that 92.2% of all cells were fibroblasts, while spatial transcriptomics analyses indicated that 68.1% of all cells were fibroblasts (Figure 2A and S9a). Despite these differences, both methods demonstrated a high level of concordance between the cell type populations identified (Figure 2a and S9a), which allowed for an integrated further analysis. Spatial transcriptomic unsupervised clustering further enabled the identification of distinct keratinocyte subclusters, including inner root sheath (IRS) keratinocytes (positive for KRT71), bulge region keratinocytes (enriched for LHX2), and keratinocytes of the matrix (positive for MKI67). Finally, few KRT20+/ATHO1+ Merkel cells were detected by spatial transcriptomic in close proximity to the hair follicles (Figure 2c-d). Furthermore, we detected the presence of several clusters of dying cells characterized by the expression of KRT10 and primarily localized in the core of the organoids. These cells are likely to represent squamous cells, which accumulate in the center of the organoids (Figure S9). Due to the limited number of markers in spatial sequencing, as well as the thinning that occurs in the stratified epithelium at high days of differentiation, outer root sheath (ORS) and basal layer cell types (both sharing positivity for KRT17 and KRT5) as well as stratum spinosum and granulosum (positive for GATA3 and MALL, respectively) could not be precisely distinguished and instead grouped together in one cluster each (Figure 2c and Supplementary Figure S9). The fibroblast population was re-analyzed in the scRNA-seq data by applying known expression patterns of different fibroblast subpopulations previously identified in adult skin human samples ^30–32^. Consequently, a mesenchymal fibroblast cluster was identified on the basis of the expression of ASPN, SFRP1, and IGF1 ^31^. Furthermore, according to the role of mesenchymal fibroblasts in supporting the deposition of cartilage and bone marrow ^32^, mesenchymal fibroblasts were clearly localized around the tail of the organoids, where chondrocytes and cartilage are found (Figure 2h-i and Supplementary Figure S9b). Papillary fibroblasts have been shown to localize within the first 100µm layer below the stratified epithelium, and their differentiation strictly depends on the keratinocytes in their vicinity ^30^. The papillary signature was characterized by the positivity of PTGDS, MEF2C, and RSPO1 ^30,31^, and these cells were found within the epithelium and the outer limit of the SkOs (Figure 2e-h and supplementary Figure S9b-c). The analysis of marker expression and localization has enabled the further subdivision of papillary fibroblasts into three distinct subtypes sharing many markers (Figure 2h-i and Supplementary Figure S9c). However, while papillary fibroblast subtype 1 is represented by a population of fibroblasts in the outermost layer of the organoid with higher levels of THY1 and PTGDS markers, papillary subtype 2 fibroblasts, in close contact with the basal layer of the skin, are characterized by higher levels of MEF2C. Finally, papillary fibroblast type 3 are mostly single cells distributed in the papillary region.

**Fig. 2.**
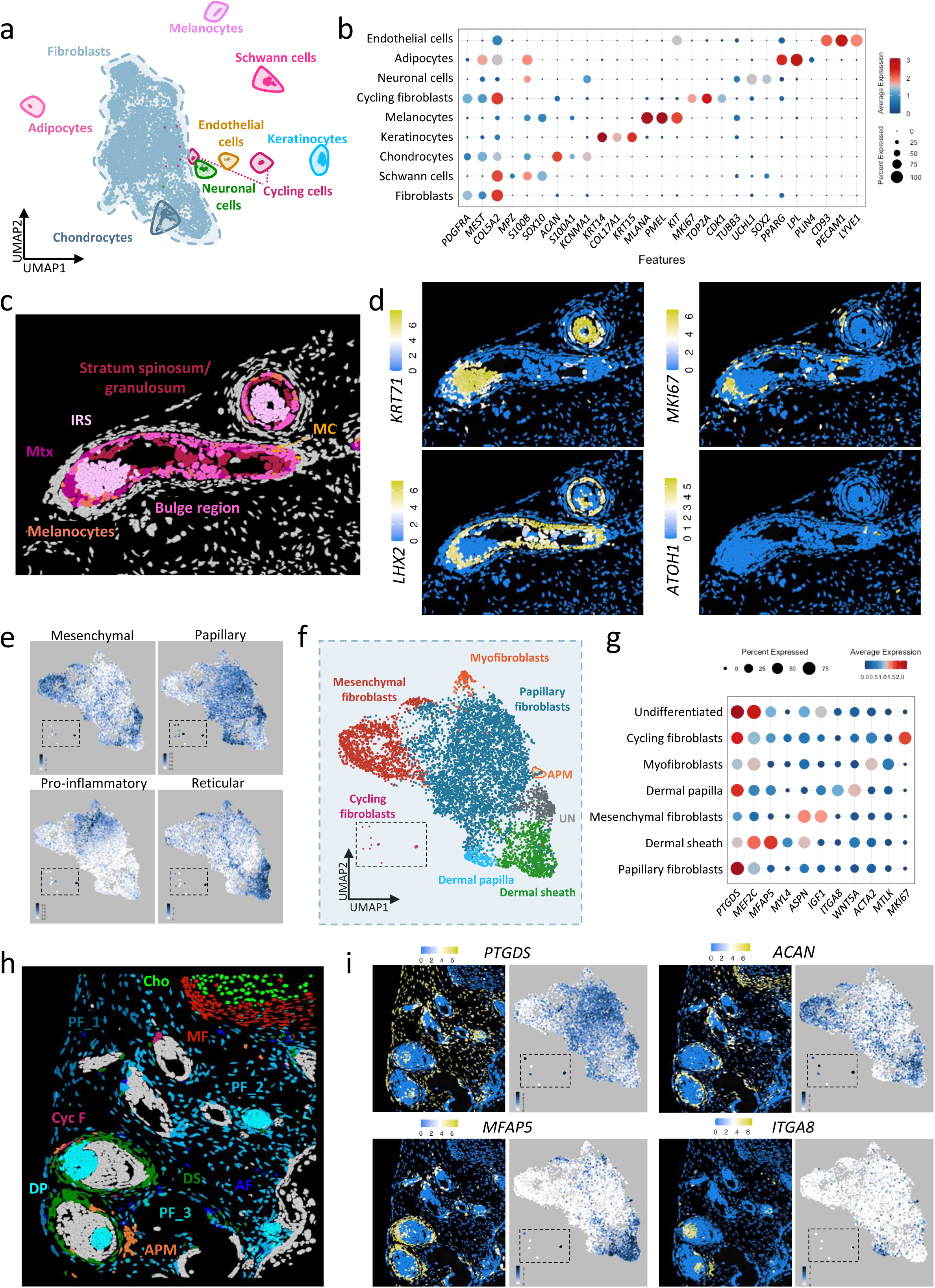
Combination of scRNAseq and spatial transcriptomics define the different cell type populations and localization in the SkOs. a, Uniform manifold approximation projection (UMAP) plot of 150-day SkOs. Each point represents a single cell (7,658 cells in total from non-infected and 100- and 120-infected SkOs). The colors indicate the different clusters defined based on the expression of cell type-specific markers. b, Dot plot showing the expression of cell type-specific markers across the different cell clusters. The size of the icons represents the percentage of positive cells; colors code for average expression. c, Spatial visualization of a region of interest (ROI) from a non-infected organoid showing the localization of different keratinocyte populations in the SkO. d, Spatial feature plots of the ROI in panel c, colored by expression level of the indicated genes. e-g, The fibroblast population from panel a was subset and reanalyzed using unsupervised clustering to identify different fibroblast subpopulations. Dashed boxes show the subset of cycling fibroblasts from panel a. e, Average expression of the genes constituting the mesenchymal (*APSN* and *SFRP1*), papillary (*APCDD1, COL18A1, WIF1,* and *PTGDS*), pro-inflammatory (*CCL19*, *APOE*, *CXCL2*, *CXCL3*, and *EFEMP1*), and reticular (*MFAP5, POSTN* and *SLPI*) fibroblast gene signatures. f, UMAP plot of the fibroblast and cycling fibroblast subsets. Colors indicate the different clusters defined based on the combination of the fibroblast-specific marker expression and spatial information. g, Dot plot showing the expression of fibroblast subtype-specific markers across the different cell clusters. The size of the icons represents the percentage of positive cells; colors code refers to the average expression. h, Spatial visualization of a non-infected SkO ROI showing the distribution of the different fibroblast subpopulations. Cho: chondrocytes, MF: mesenchymal fibroblasts, Cyc F: cycling fibroblasts, DP: dermal papilla, APM: arrector pili muscle, PF: papillary fibroblasts, AF: activated fibroblasts. i, For the indicated genes, the spatial feature plot of the ROI in panel h, and the feature plot of the scRNA-seq fibroblast and cycling cell subset are shown.

Based on a proinflammatory signature (depicted as average expression levels, Figure 2e) characterized by the expression of CCL19, APOE, CXCL2, EFEMP1 and CXCL3 ^31^, we identified a further subpopulation of papillary fibroblasts in single-cell analysis, which we cannot trace in spatial transcriptomics due to limitations in the number of probes. We occasionally find clusters of activated papillary fibroblasts (negative for APOE, a marker for pro-inflammatory fibroblasts, and enriched for ISG15 and IL6) in the vicinity of necrotic cells (Supplementary Figure S9c).

In adult skin, reticular fibroblasts are located in the layer beneath papillary fibroblasts (at a depth of 100– 300 µm) and are characterized by the expression of MFAP5, SLPI, CTHRC1 and TSPAN^31^. The marker MFAP5, a hallmark of reticular fibroblasts, exhibited a characteristic expression pattern in the scRNA-seq data that was consistent with cells localized near hair follicles, where the dermal sheath is expected (Figures 2e-i). Furthermore, high levels of DPEP1 and MYL4 ^33^, known markers for the dermal sheath population, are also highly enriched in this population. Consequently, reticular fibroblasts are thought to be absent from the SkO due to the configuration of the organoids, and this cell population was therefore designated the dermal sheath. Finally, in scRNA-seq, we detected a population of cells characterized by the expression of markers for papillary, mesenchymal, and reticular fibroblasts (here named undifferentiated, Figure 2e-g) that did not appear in the xenium clustering. Consequently, we hypothesize that these cells were randomly assigned to the mesenchymal, papillary, and/or reticular clusters in the spatial transcriptomic analysis.

Spatial transcriptomic analysis was fundamental to define the remaining population of cells. A small population of muscle cells, distinguished by ITGA8 expression ^32^, was identified as arrector pili muscle and found in close proximity to hair follicles (Figures 2h-i and supplementary Figure S9). These cells are only occasionally found in the SkOs and correspond to a small number of cells in the single cell analysis (Figures 2f, h and i). The dermal papilla population was distinguished by its clear localization at the base of the hair follicle and its positivity for SOX2, WNT5A, LEF1, WIF1 markers in both sequencing analyses^33^. This population also shared positivity for ITGA8, a known marker of arrector pili muscle (Figure 2i).

In conclusion, given the paucity of sequencing information on end-stage differentiated SkOs and their differences from human adult skin, a combination of scRNA-seq and spatial transcriptomic analyses was fundamental to determine the different cell populations of the organoid.

### MCPyV preferentially infects papillary and dermal sheath fibroblasts in the SkO

Given that productive MCPyV infection could only be observed in the context of the organoid, we reasoned that a particular state of differentiation or subpopulation of fibroblasts is necessary to support the full life cycle of the virus. We therefore used various methods to examine the viral genome, viral RNA, and viral protein expression by DNAScope, RNAScope for early and late gene expression, and immunohistochemistry, respectively. Consistent with our observations in WMI with VP1 expression, immunohistochemistry revealed a substantial number of cells with LT expression (Figure 3a, upper panel). Remarkably, the detection of the viral genome indicated that the majority of dermal fibroblasts carry the genome, though only a subset of these expressed the viral protein (Figure 3a, lower panel and Supplementary Figure S4). A more thorough examination of viral transcription using RNAScope corroborates the IHC and WMI results, with only a subset of cells manifesting viral transcripts (Figure 3b) and viral protein expression (Figures 1a and 3a, upper panel, and Supplementary Figure S2 and S3), while the genome is present in nearly all cells (Figure 3a, lower panel and Supplementary Figure S4). The employment of a combination of distinct RNA probes in RNAScope, individually detecting sT/LT and VP1, facilitated the detection of cells in various phases of infection (Figure 3b). Specifically, the analysis revealed cells that exclusively expressed early transcripts (Figure 3b, panel 1), cells exhibiting comparable expression of early and late transcripts (Figure 3b, panel 2), and cells with a high number of late transcripts, which are likely to be productive cells of virus progeny (Figure 3b, panel 3). These results were also identified in the scRNA-seq data (Figures 3c-e), which revealed diverse infected cell populations exhibiting varying ratios of early and late viral gene expression, as would be expected in an authentic infection (Figure 3e).

**Fig. 3.**
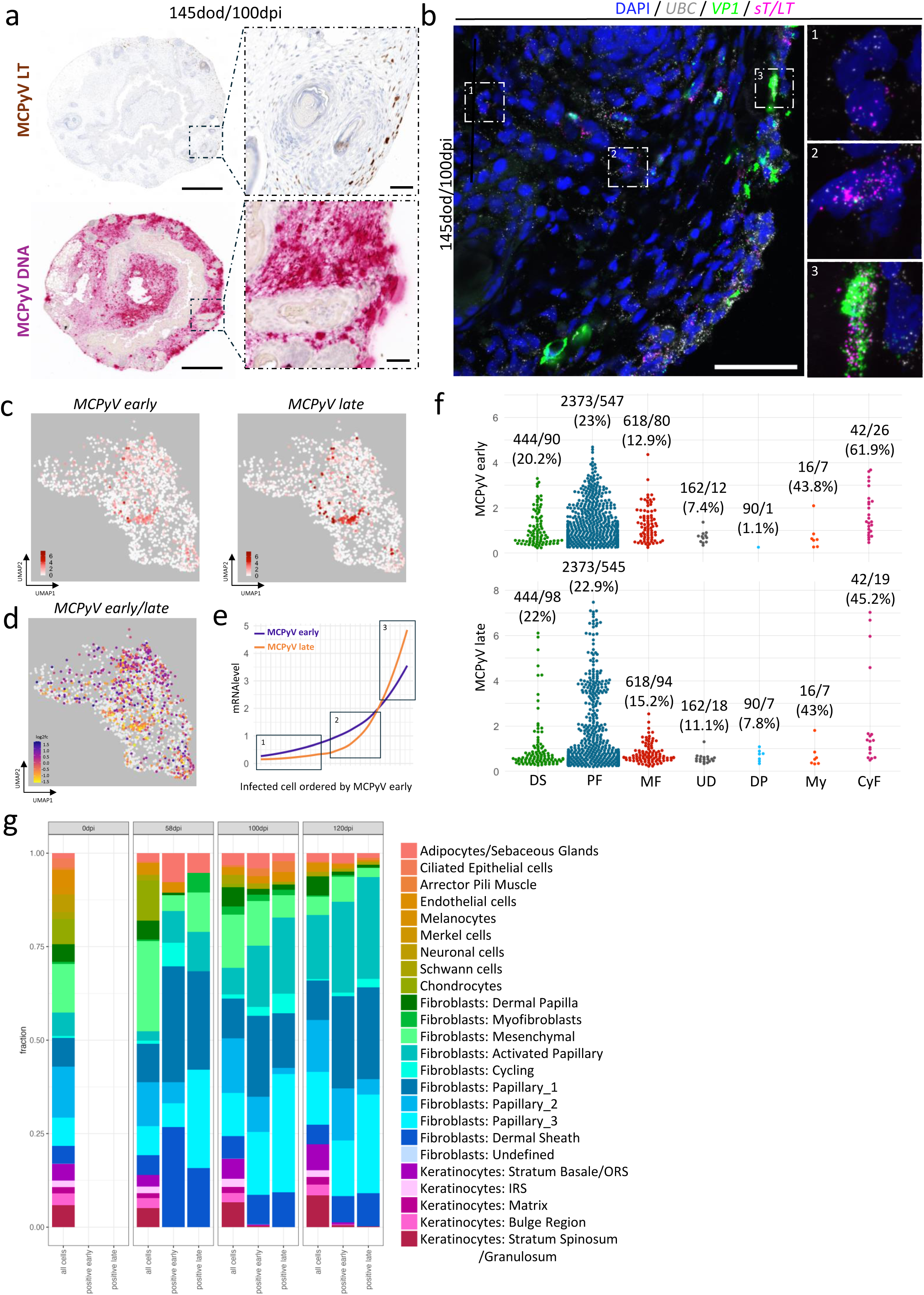
MCPyV early and late gene expression is preferentially supported by papillary fibroblasts in the SkOs. **a,** 145-day-old SkOs infected for 100 days (145dod/100dpi) were formal-infixed paraffin-embedded (FFPE) and serial sections were labelled by immunohistochemistry for MCPyV LT (Ab3) antigen (top), and in situ hybridization DNAScope (red) visualizing the MCPyV DNA (bottom). Dashed boxes indicate the magnified regions on the right. Scale bar: 500 μm for the main images, 50 μm for the magnifications. **b,** In situ RNAScope of fresh-frozen 145dod/100dpi SkOs showing expression of MCPyV sT/LT mRNA (magenta), MCPyV VP1 mRNA (green), ubiquitin C (UBC, housekeeping, grey); nuclei are counterstained with DAPI. Dashed boxes and numbers indicate the magnified regions to the right, with representative examples of (1) cells expressing only viral early transcripts, (2) cells expressing both early and late transcripts, with higher levels of early transcripts, and (3) cells expressing predominantly late transcripts. Scale bar: 200 μm. **c,** Feature plot of MCPyV early (left) and late (right) mRNA expression across single cells in the fibroblast subclusters. Each point represents a single cell, and the colour scale corresponds to the normalized expression level, with darker colours indicating higher expression levels. **d,** Feature plot showing the log2 fold change (log2fc) between MCPyV early and late mRNA expression across single cells. Colour indicates the direction and magnitude of the fold change: log2fc > 0 (blue) indicates cells in which viral early gene expression is higher than late gene expression, while log2fc < 0 (yellow) indicates cells in which viral late gene expression is higher than early gene expression. **e,** Smoothed expression profiles of MCPyV early and late genes across single cells, ordered along the x-axis based on the expression level of MCPyV early mRNA. The level of early (blue) and late (orange) MCPyV mRNA are shown as trend lines. Black insets indicate the different stages of MCPyV infection, as in panel b. **f,** MCPyV early (top) and late (bottom) gene expression across fibroblast cell clusters. The total number of cells/number of infected cells is shown for each cluster. The percentage of infected cells is given in parentheses. DS: dermal sheath, PF: papillary fibroblasts, MF mesenchymal cells, UD: undifferentiated, DP: dermal papilla, My: myofibroblasts, CyF: cycling fibroblasts,. **g,** Stacked bar plot showing the percentage of all, early infected, and late infected cells per cluster in each condition.

A comprehensive analysis of early and late transcripts in our single-cell RNA sequencing (scRNA-seq) dataset confirmed that fibroblasts constitute the predominant cell type in the infected sample, accounting for 97.8% of the infected cell population (Figure 3f and Supplementary Figure S10). In addition, a smaller population of cycling fibroblasts expressing viral transcripts was identified. A comparative analysis of early and late viral transcripts in distinct fibroblast subpopulations reveals substantial differences among the subtypes, particularly with respect to the expression levels of late viral transcripts. In this regard, papillary fibroblasts, and to a lesser extent, dermal sheath fibroblasts, exhibit a cluster of cells that demonstrate a significantly higher level of late viral transcript expression compared to other fibroblasts, except for a few cycling cells (Figure 3f). Interestingly, papillary and to a lesser extend dermal sheath and cycling fibroblasts each show a cell population within the clusters with increased levels of late transcripts (Figure 3f). Spatial analysis yielded analogous results, with papillary fibroblasts (Figure 3g) comprising 95% of infected cells. Remarkably, papillary fibroblasts constituted the majority of infected cells in both, the early and late stages of infection (Figure 3g). Interestingly, we find an increased population of activated fibroblasts in infected organoids, which exhibited a substantial enrichment of IL6 and ISG markers in the infected area, (Figure 3g, turquoise population). This observation suggests that MCPyV infection stimulates the activation of papillary fibroblasts.

In summary, the results indicate that MCPyV infection in the skin organoids is predominantly restricted to papillary and dermal sheath fibroblasts and preferentially establishes viral late gene expression in papillary fibroblasts in the infection model employed.

### MCPyV subverts the host immune response in SkO

In order to explore the modulation of host gene expression in the SkOs infected with MCPyV, a transcriptomic analysis was performed in bulk of pools of non-infected (CTRL) and 293-MCPyV infected SkO pools at 100dpi and 120dpi (Supplementary Figure S7). The analysis of the deregulated genes (DEG) at both time points vs non-infected SkOs showed a strong upregulation of interferon-stimulated genes (ISGs) with GO term analysis confirming the upregulation of pathways involved in response against infection and innate immune response (Figure 4 b and Supplementary Figure S11b-c). The increase in the expression of ISGs in MCPyV-infected organoids compared to control SkOs was also detected on a single cell level, e.g. scRNA-seq and spatial transcriptome analysis (Figure 4c-f and Supplementary S11).

**Fig. 4.**
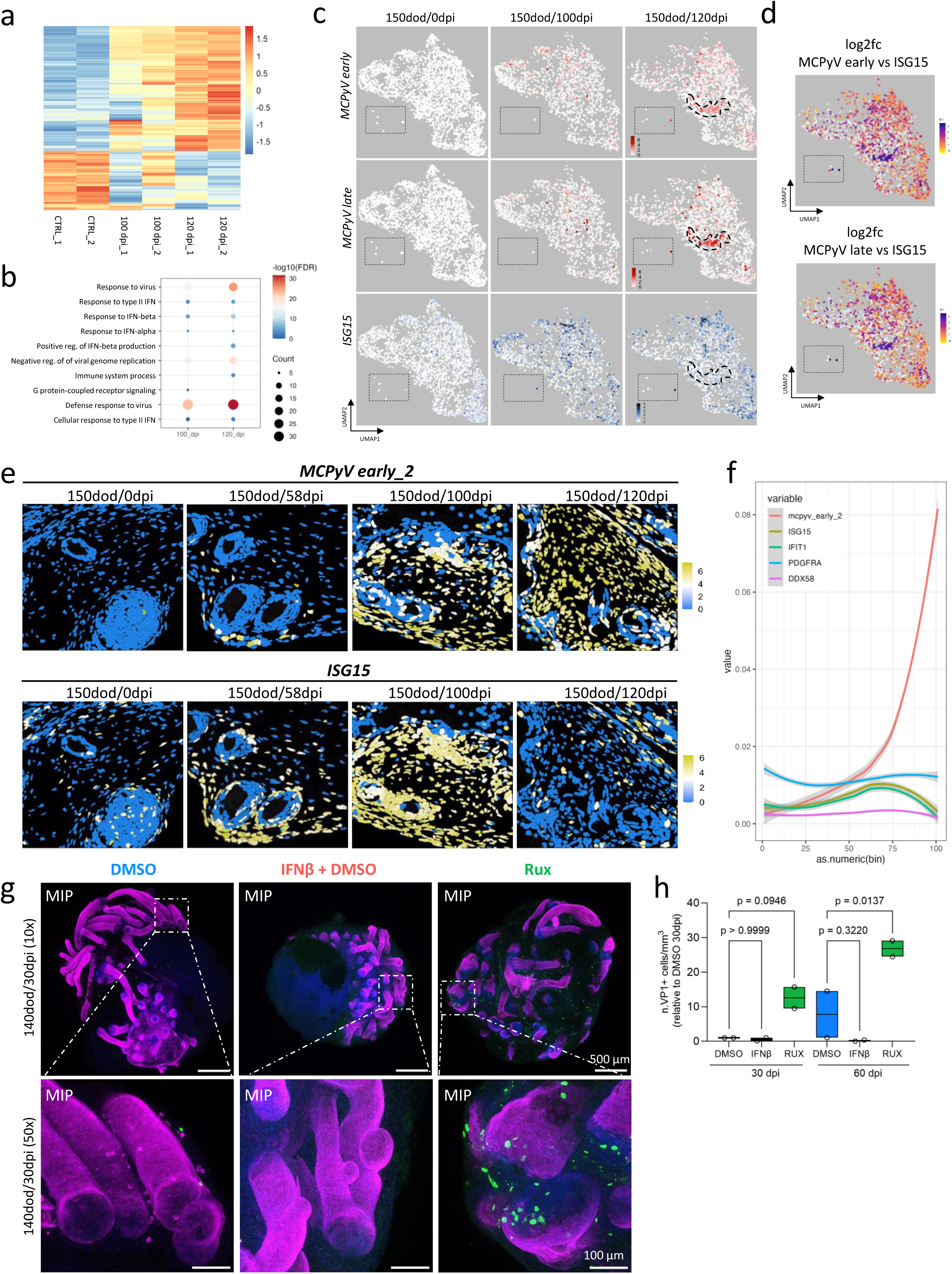
MCPyV subverts ISG production to establish a productive infection in papillary fibroblasts. **a,** Heat map of deregulated genes (DEGs) obtained from SkOs infected for 100d and 120d and analysed against non-infected organoids (CTRL). The colour code refers to the row Z-scores. **b,** Gene Ontology (GO) analysis (DAVID Bioinformatics Resources) was performed using significantly (padj. ≤0.05) upregulated (log2FC ≥1) genes from the RNA-seq analysis. From all significantly enriched GO terms with a minimum gene count of five, the ten terms with the highest gene counts were derived. The size of the icons represents the gene counts; the colour code refers to the level of significance (FDR). **c,** Feature plot for the fibroblast and cycling fibroblast cells (dotted insets) subsets showing the gene expression of the indicated genes in each cell. The dotted line indicates a group of highly infected cells. **d** Feature plot showing the log2 fold change (log2fc) between MCPyV early (top) or late (bottom) and *ISG15* mRNA expression across single cells. Colour indicates the direction and magnitude of the fold change: log2fc > 0 (blue) indicates cells in which viral early gene expression is higher than *ISG15* gene expression, while log2fc < 0 (yellow) indicates cells in which viral gene expression is lower than *ISG15* gene expression. **e,** Spatial visualization of a region of interest (ROI) of non-infected (150dod/0dpi) and infected SkOs (150dod/58dpi, 150dod/100dpi, 150dod/120dpi, respectively). The color code refers to the expression level of the indicated gene. **f,** Cells with at least four viral counts in the spatial analysis were ordered along the viral load and binned with 20 cells per bin. The percentage of MCPyV early transcripts and *ISGs* within the bin is shown. *PDGFRα* is used as internal control. **g,** Whole mount immunostaining of 140dod/30dpi and treated with vehicle control (DMSO), IFNβ or ruxolitinib (RUX), throughout the infection. KRT17 staining for epidermis and outer root sheath of the hairs (magenta), MCPyV VP1 staining for the virus capsid protein (green), and Hoechst counterstaining for nuclei (blue). The images show the maximum intensity projection (MIP) of multiple z-stack acquisitions. Dashed boxes indicate the magnified regions. Scale bar: 500 μm in the main images, 100 μm in the magnified images. **h,** For each sample, the number of VP1 positive cells in the SkO was quantified and normalized to the volume. Shown is the mean and standard deviation of two independent experiments with a total of five biological replicates per condition and normalized to DMSO 30dpi. One-way ANOVA followed by Sidak post hoc test.

It is interesting to note that in the population of papillary fibroblasts, a group of cells in which there is a high level of early and late MCPyV gene expression, shows a significant reduction in ISG expression (Figure 4c-d). Furthermore, spatial analysis revealed an increase in ISGs in both infected cells and their surrounding bystander cells, suggesting a paracrine effect of infection on neighboring cells. ISG levels appeared to increase progressively with the percentage of infected cells and then decrease in areas of maximal infection (Figure 4e-f).

Following an initial stimulation of the innate immune response after MCPyV infection, we hypothesize that the virus must subvert the immune response to establish an efficient and productive infection. To test this hypothesis, we challenged the infection by IFNβ treatment, to boost the expression of ISGs, as well as with Ruxolitinib (RUX), a JAK inhibitor, to block ISG production ^34^. We then assessed the number of VP1-positive cells at 30 and 60dpi by WMI. Similar to previous results in infected fibroblasts grown as a monolayer, we find MCPyV infection to be significantly reduced by IFNβ treatment (Figure 4g-h and Supplementary Figure S12), with VP1 positive cells only rarely observed even after 60 dpi. Conversely, while RUX did not increase the infection rate in monolayer culture^34^, an upregulation in the number of VP1-positive cells was observed at 30 and 60dpi in the SkOs (Figure 4g-h and Supplementary Figure S12).

Collectively, the results strongly suggest that MCPyV infection initially stimulates the innate immune response in SkOs. However, the virus must subvert the host immune response to efficiently reactivate and establish a productive and efficient infection in the SkOs. Furthermore, this effect appears to be more efficient in papillary fibroblasts, where high levels of early and late viral gene expression can be observed in cells with reduced levels of ISGs.

## Discussion

MCPyV, like all human polyomaviruses, is a virus that has co-evolved with and is highly adapted to humans, which makes the development of in vitro infection models challenging. This is evidenced by the lack of simple infection models for human polyomaviruses and, in the case of MCPyV, the lack of infection systems in general that result in the production of viral progeny. Further, there are no small animal models available to study MCPyV infection; current animal models based on transgenic mice are highly valuable in depicting tumor progression ^15–17,35,36^. Existing in vitro cell culture models to study some aspects of MCPyV life cycle either utilize circularized viral genomes that are transfected into cell lines or primary cells or generate low levels of MCPyV particles in helper cells that are subsequently employed for infection. However, all these systems do not result in the production of viral progeny and reinfection. The issue of the paucity of simple *in vitro* infection models for human polyomaviruses also pertains to related human papillomaviruses (HPV). However, the majority of HPV subtypes can be studied in raft cultures, which model a stratified epithelium. Keratinocyte differentiation in these cultures enables HPV replication and release. However, it is important to note that raft cultures do not support the life cycle of MCPyV.

The knowledge of the cells and tissues in which MCPyV establishes and maintains its infection is further based on the semi-permissive models described earlier together with epidemiological studies. MCPyV is a highly prevalent virus that can be detected in skin swabs from individuals using highly sensitive PCR, suggesting that the skin is the site of virus persistence ^37^. In addition, studies have identified the presence of the virus in the hair bulge of plucked eyebrows, suggesting its ability to persist within the cells of hair follicles. Also, the vast majority of tumors induced by MCPyV, MCC are located in the dermis, close to the hair follicles ^38^.

One in vitro infection model for MCPyV is particularly noteworthy: an in vitro infection model in skin fibroblasts that allows initial infection and mimics parts of the viral life cycle, but does not lead to the release of viruses ^11^. In this model, studies were also carried out on the role of the innate immune response, which demonstrated robust activation of the ISG response after MCPyV infection and the release of inflammatory cytokines, such as IL-1b, IL-6, TNF-a, and IL-8 ^14,34^. These results were also confirmed by overexpression experiments ^13^. However, it should be noted that MCPyV infection can only be followed for a limited period of up to 5-7 days in all systems used so far, without evidence of virus production and spread in the culture ^11,13^. With regard to other viruses, such as papillomaviruses and VZV, the differentiation of the skin may also be of crucial importance for MCPyV. However, in the case of polyomaviruses, it is possible that different subpopulations or differently differentiated fibroblasts could be of crucial importance. Nevertheless, since fibroblasts lose their differentiation relatively quickly in vitro, this could at least partially explain the divergent reports of infection rates of 1-5% and 20-40%^11,13^.

Based on all these indications, we have explored a recently published skin organoid model ^20,21^ as a putative infection model for MCPyV. Skin organoids, derived from iPSCs, constitute a complex tissue comprising a diverse array of skin cell types and multilayer structures. Notably, this model has recently been employed in the context of acute, short-term (up to 7d) infections involving skin-rooted viruses, such as SARS-CoV-2 and MPoX ^18,19^.

As with many complex organoid models, SkOs also exhibit an inside-out morphology, which poses two challenges: firstly, the route of infection and, secondly, the limited lifetime of the organoids, with epithelial cells released to the lumen of the organoid and necrosis setting in after 150 days of differentiation. The exposure of the dermis to the external environment is actually advantageous for MCPyV infection, given that fibroblasts are important for infection. However, the limited lifespan of the organoids poses a challenge, particularly for a slow virus such as MCPyV, insofar as there might be insufficient time to reach cell layers other than the dermis. Indeed, we observe a slow infection in the case of MCPyV, requiring approximately 100 days to achieve significant amount of infected cells expressing the viral VP1 protein. Another challenge is the relative rigidity of the organoids at later points in differentiation, to which the extracellular matrix provides a barrier to virus entry into the organoid, as we have shown in DNAScope experiments. When infecting early in differentiation and following for extended time periods, the various stages of the virus life cycle can be visualized in dermal fibroblasts using different methodologies including transmission electron microscopy (TEM). Furthermore, we demonstrate successful virus production and re-infection for the first time with MCPyV produced in the SkOs being morphologically and genetically identical to the original virus without evident rearrangements in the virus’s promoter region, the NCCR, as described for some human polyomaviruses e.g. BKPyV ^39^.

The presence of the viral genome in the plethora of fibroblasts of the organoid was an unexpected finding. In contrast to the detection of viral transcripts and viral protein expression, a significantly higher number of fibroblasts were found to carry the viral genome. This suggests that MCPyV can enter most fibroblasts, but only a fraction of them support viral progeny production. This is a particularly interesting finding considering our previous finding of long-term persistence of MCPyV genomes in a cell culture system^22^. This investigation employed 2D cell culture systems, that had been transfected with MCPyV genomes. The resulting cultures were observed over several months with the copy number of the viral genomes remaining stable even beyond several months. At that time, together with our findings that an MCPyV genome mutant that does not express its viral miRNA and therefore has higher levels of LT protein is lost significantly faster, we speculated that LT expression, which is known to be highly immunogenic, and its tight regulation contribute to viral genome clearance. This study’s long-term infection experiments in SKOs over several months support the hypothesis that MCPyV establishes a distinct form of persistence that has not been previously described for polyomaviruses. Subsequent experiments in follow-up studies will investigate the precise mechanism of how this persistence is regulated.

Interestingly, another viral protein, ALTO has been proposed recently to contribute to persistence ^40–43^. While the role of ALTO in the viral life cycle is unclear, recent work suggests that ALTO has a role as a tumor suppressor, which is inactivated in MCC ^42^. Interestingly, ALTO regulates early viral protein expression via NF-kB activation and binding to the NCCR of MCPyV genomes ^41,42^. This suggests a mechanism by which ALTO, regulates the expression of early viral proteins, thereby ensuring that LT expression is tightly controlled indicating a delicate balance between viral proteins and mechanisms and the cellular innate immunity. In previous studies, it has been demonstrated that infection with MCPyV in dermal fibroblasts results in the activation of STING, TBK1, IRF3 and NF-kB, subsequently inducing an ISG response ^13,14,34,42^. This observation was successfully replicated in our model, where the ISG response was found to be significantly induced by MCPyV infection. However, our single-cell and spatial transcriptome analyses demonstrate that the virus can evade this ISG response in certain cell types, resulting in an inverse correlation between ISG expression and early MCPyV gene expression. This observation is of particular interest, as we have recently demonstrated in primary cells transduced with MCPyV sT that sT is capable of circumventing innate immunity through the reduction of the ISG response mediated by IRF9 and ISGF3^13^. The investigation of the axis of innate immunity regulation in the viral context in SKOs is of significant interest.

In the present study, a combination of methodologies was employed, encompassing longitudinal single-cell transcription analyses of infected organoids in conjunction with spatial transcriptome analyses conducted at three distinct time points of infection. The integration of these approaches enabled the delineation of cell types based on their expression patterns and the determination of their precise location within the tissue, with a high degree of resolution at the fibroblast level. Since there is little sequencing information available on human skin organoids at a later stage of differentiation, we used an existing set of probes in the Xenium-based probe set that reflect the expression of specific markers of adult skin, and expanded it with 100 custom probes that we designed based on our previous results to target the expression of innate immunity genes and viral transcripts. This has its limitations, given that the SkO model mimics human second-trimester fetal skin ^20^, and due to the differentiation of iPSCs, we may not have fully differentiated progenitor cells in the organoid. The interpretation of results and their careful consideration necessitates validation through alternative techniques, such as single-cell RNA sequencing (scRNASeq), RNAScope analysis, and immunohistochemical staining.

Using a combination of high-resolution methods, we were able to show that MCPyV primarily infects dermal fibroblasts. We were able to show that certain fibroblast subtypes, papillary fibroblasts and dermal sheath fibroblasts, show productive MCPyV infection. Subsequent transcriptome analyses showed that MCPyV only expresses elevated levels of early and late viral transcripts mainly in papillary fibroblasts and, interestingly, the expression levels of key ISGs are reversed, effectively evading the cellular immune response to the virus. The possibility that mesenchymal and dermal sheath fibroblasts, which are located further inside the organoid, reactivate more slowly than papillary fibroblasts cannot be ruled out at present. The identification of dermal cells that settle in the hair root layers as the infectious cell type is of considerable relevance for several reasons. Firstly, MCPyV was detected in the hair follicles of plucked eyebrows using PCR. Secondly, these cells also represent an important first cell population for infection with regard to the infection pathway through the hair canals. This is a process that is undoubtedly regulated by the immunological niche in the skin in this area ^44^, which consists of macrophages and Langerhans cells, and the virus is rapidly contained. In subsequent studies, we will introduce immune cells, specifically macrophages, into the model and further investigate this hypothesis.

The model is innovative in several ways for the field of MCPyV and viral tumorigenesis in general: in mapping and understanding the infection cycle, in understanding a novel form of viral persistence, in further developing it as a pathogenicity model, and also in testing new approaches to intervention against MCPyV reactivation, a prerequisite for tumorigenesis.

## Material and methods

### Skin organoids differentiation

Skin organoids (SkOs) were generated using UKEi001-A (RRID:CVCL_A8PR) human induced pluripotent stem cells isolated from a female adult and used between passages 31-39. All procedures involving the hiPSC lines were approved by the local ethics committee in Hamburg (Az PV4798, 28.10.2014). IPSCs were cultured in 6-well plates in StemFlex medium (SFM, Gibco) on gel Trex-coated plates (Gibco). SkO generation was performed as previously described ^20,21^, with some modifications. Briefly, 3,000 iPS cells were plated in 100 μl Essential E8 (Gibco) medium in a 96-well U-bottom plate (Nunclon Sphera) with 100 ug ml^-1^ of Normocin (Invivogen) and 10 μM Y27632 (Biorbyt). At day 0 of differentiation, the concentration of BMP4 was adjusted to 2-3.5 ng ml^-1^ depending on the size of the spheroids. After day 12, the SkOs were cultured in 500 μl of organoid maturation medium (OMM) in a 24-well plate. At day 70, the volume of the OMM medium was increased to 1 ml.

### MCPyV virus production, infection and treatment

Lenti-X 293T-derived MCPyV virions (293-MCPyV) were generated as previously described ^45^, with minor modifications. Lenti-X-293T cells (ATCC) were transfected with pMtB (Addgene #32096), pALD* (Addgene #32097), and the re-ligated recombinant genome of MCPyV ^12^ using Lipofectamin 2000 (Invitrogen). MCPyV virions were harvested 9 days post-transfection and purified with an OptiPrep gradient. Fractions were screened by Western blot for VP1 (Christopher B. Buck, NCI) and silver staining. Positive fractions were dialyzed using dialysis cassette (Spectra/Por). Viral titers were calculated as genome equivalents by quantitative PCR.

Infection of nHDFs (Lonza, #CC-2509) was performed as previously described^11,13^ with 10E5 viral particles/cell. For SkO infection, 10E8 293-MCPyV was added to each organoid in 500 μL of fresh OMM medium. If infection was performed in organoids <70 days old, 500 μL OMM was added to the well the following day to reach a final volume of 1 ml. Medium was then changed every other day according to the SkO generation protocol, with one complete medium change per week. For IFNβ (PeproTech) and Ruxolitinib (RUX, ChemCruz) treatment, 5ng/mL and 5µM of the compound, respectively, were added to each well 2 days prior to infection and maintained until organoid fixation, added with each medium change.

### Immunostaining

Whole-mount immunostaining of organoids was performed as previously described^20,21^ following the qCd3e protocol. SkOs were mounted in clearing solution between two glass coverslips, which were spaced by 1 mm spacers (SunJin Lab). Each organoid was scanned from the top and the bottom view using a confocal laser scanning microscope (DMi8, Leica TCS SP8 X, Leica). To quantify the number of VP1 positive cells per organoid, the overlapping stacks between the top and bottom view were manually removed using the FIJI software. The number of positive cells was calculated as the sum of the top and bottom views normalized to the SkO volume, which was calculated using the IMARIS 9 software.

Cells were fixed with 4% paraformaldehyde in PBS for 10 min to perform immunofluorescence on MCPyV infected cells. Staining was performed as previously described ^13^. Images were acquired using a confocal laser scanning microscope (DMi8, Leica TCS SP8 X, Leica).

FFPE immunohistochemistry and immunofluorescence were performed on organoids which were fixed in a Leica ASP300S tissue processor and embedded in paraffin. Paraffin sections of 3 μm were cut and stained with hematoxylin and eosin (H&E) according to standard procedures.

Organoid sections were processed for immunohistochemical staining as follows: After deparaffinization and inactivation of endogenous peroxidases (3% hydrogen peroxide), antibody-specific antigen retrieval was performed using the Ventana BenchMark XT autostainer (Ventana). Sections were blocked and incubated with the primary antibodies. The UltraView Universal DAB Detection Kit (Roche, #760-500) containing both, anti-mouse and anti-rabbit secondary antibodies was used for the detection of specific binding and DAB staining. Counterstaining and bluing were performed with Hematoxylin (Ventana Roche, # 760-2021) and Bluing Reagent (Ventana Roche, # 760-2037) for 4 minutes, followed by the mounting of stained sections in mounting medium.

For immunofluorescence staining, paraffin organoid sections were deparaffinized thoroughly in xylene and a descending alcohol series for 2x 20 minutes. Antigen retrieval was then performed by pressure boiling the sections in Universal R buffer (#AP0530-500; Aptum) for 20 minutes. Sections were rinsed briefly and blocked for 1 hour. Primary antibodies were incubated overnight at 4°C. After intensive washing, secondary antibodies conjugated to AlexaFluor488-, AlexaFluor555, or AlexaFluor647 were applied for 1.5 hours. Sections were washed, counterstained with DAPI and mounted in Fluoromount-G (SouthernBiotech). Data acquisition was performed using a Leica Sp8 confocal microscope and Leica application suite software (LAS-AF-lite). All the antibodies used are listed in Table S1.

### DNA in situ hybridization

MCPyV DNA was detected on 4 µm FFPE sections of infected organoids using the DNAScope™ Duplex Assay Kit (Advanced Cell Diagnostics) and a full genome probe (Cat No. 1088231). The manufactureŕs instructions were strictly followed, with the addition of protease digestion using Protease Plus at 40°C for 15 minutes and target retrieval at approximately 99°C for 20 minutes. The signal was visualized using an alkaline phosphatase catalyzed chromogenic substrate precipitation. Images were acquired using a Pannoramic MIDI II scanner (Sysmex).

### RNA in situ hybridization

MCPyV mRNA was detected on 12-µm cryostat sections of infected organoids using the RNAScope™ Multiplex Fluorescent V1 assay kit (Advanced Cell Diagnostics) and the following probe combination: large T/ small T (Cat No. 431001), VP1 (Cat No. 1312631-C3), and ubiquitin C (UBC) as a housekeeping gene to control for RNA integrity (Cat No. 310041-C2). Signals were visualized using Atto Fluor 550, Atto Fluor 647, and Alexa Fluor 488, respectively. To validate the MCPyV signal in the Xenium experiments, adjacent 4-µm FFPE sections were stained with the same probe combination using the RNAScope™ Multiplex Fluorescent V2 assay kit (ACD) and standard pretreatment conditions (30 min protease plus at 40°C and 15 min target retrieval at approximately 99°C). Signal was visualized using the TSA Vivid fluorophores 630, 520 or 570 (Tocris Bioscience), respectively. Images were acquired using a confocal Olympus FV3000 microscope (IX83, Olympus).

### Quantitative polymerase chain reaction

Quantitative polymerase chain reaction (qPCR) was performed to detect MCPyV genome copies in the supernatant of infected SkOs using VP1-targeting primers (5’-TAAAGGAGGAGTGGAAGT-3’ and 5’-CATTTAGCATTGGCAGAGA-3’) and a fluorescein amidite (FAM) labelled TaqMan® probe (5’-[6FAM]GATCTGGAGATGATCCCTTTGGCTG[BHQ1]-3’). The supernatant was heat-inactivated at 95°C for 10 min and diluted 1:5 in distilled DNase/RNase-free UltraPure™ water (Thermo Fisher Scientific). Reactions and analysis were performed on a LightCycler® 480 II (Roche Diagnostics) with absolute quantification/fit points analysis using 4 fit points and a threshold calculated automatically calculated by the program. A standard and a positive control of an MCPyV plasmid were used to calculate genomic copies. The supernatant of 4 to 6 organoids was analyzed for each condition. The measured MCPyV genome copies per µl of supernatant were normalized to the total amount of supernatant (500 µl or 1 ml, depending on the day of differentiation).

### Transmission electron microscopy

For negative stain transmission electron microscopy (TEM), extracellular 293-MCPyV or organoid-derived virions (SkO-MCPyV) were concentrated by ultracentrifugation to a final concentration of 2 x 10E8/µl. 4 µl of the sample was blotted onto a 300 mesh continuous carbon copper grid (Electron Microscopy Sciences), which had been glow discharged in air at 25 mA for 30 seconds in a GloQube (Quorum Technologies). The sample was negatively stained with 2% uranyl acetate and analyzed using a Talos L120C (Thermo Fisher Scientific) operating at 120 eV and equipped with a CETA camera.

Organoids were fixed overnight in 2% PFA and 2.5% GA. Subsequently, the samples were washed with PBS, postfixed for 60 minutes with 1% OsO4 in PBS, washed with PBS and ddH2O, and stained for 60 minutes with 1% uranyl acetate in water. The samples were gradually dehydrated with ethanol and embedded in Epon resin (Carl Roth, Germany) for sectioning. Ultrathin 50 nm sections were cut using an Ultracut microtome (Leica Microsystems, Germany). Sections were poststained with 1% uranyl acetate in ethanol. Electron micrographs were obtained using a 2K CCD camera (Veleta, Olympus Soft Imaging Solutions GmbH) attached to a FEI Tecnai G 20 Twin transmission electron microscope (FEI) at 80 kV. To re-locate the exact position of the TEM micrographs within the whole organoid, subsequent sections were placed on silicon wafers and examined in a Tescan Clara SEM with a BSE detector at 2.8 kV.

### Sequencing analysis

Total RNA was isolated from 4 individual organoids for each condition (uninfected, 100dpi, and 120dpi). Briefly, SkOs were first disrupted with a pestle in the QIAzol lysis reagent (Qiagen), then homogenized through a QIAshredder column (Qiagen), and finally the RNA was purified using the RNeasy MinElute Cleanup Kit (Qiagen). RNA integrity was confirmed by applying a Bioanalyzer Total RNA 6000 Nano Assay (Agilent). For each sequencing library, two independent RNA extractions from the same condition were pooled to a final amount of 1 µg total RNA (quantified by Qubit RNA HS Assay, Thermo Fisher). The pooled RNA samples were poly(A) captured using the Lexogen Poly(A) RNA Selection Kit and further processed via the RNA-Seq V2 Library Prep Kit with UDIs (Lexogen) in the long insert size variant (RTL). Libraries were quality controlled on a TapeStation D1000 Assay (Agilent, ScreenTape with D1000 Reagents) and were sequenced on an Illumina NextSeq2000 with the NextSeq 1000/2000 P2 Reagents Kit (100 Cycles) in a paired-end mode (2 x 57 bp). After demultiplexing using bcl2fastq, 21.6 – 26.3 million read pairs were assigned to each sample (pool1_non_infected: 21.6 million, pool2_non_infected: 23.7 million, pool3_100dpi: 20.3 million, pool4_100dpi: 23.4 million, pool5_120dpi: 26.3 million, pool6_120dpi: 22.1 million). Libraries from the late infection time point (120 dpi) were re-sequenced twice after the initial analysis to increase the absolute number of viral reads (2x 57 bp, NextSeq 1000/2000 P2 XLEAP-SBS Reagent Kit, 100 Cycles, and NextSeq 1000/2000 P3 XLEAP-SBS Reagent Kit, 100 Cycles, respectively, pool5_120dpi: 49.3 + 58.1 million reads, pool6_120dpi: 87.6 + 93.9 million reads). All samples passed the fastqc quality control and were subjected to downstream analysis.

### Single cell expression analysis

4 organoids for each condition (uninfected and infected for 100 and 120 days) were combined and single cell suspension was performed by mechanical grinding with a scalpel combined with enzymatic dissociation using the Tissue Dissociation Kit (Cat No. 1190062, Singleron). Organoids were incubated in dissociation buffer for 45 min at 37°C and mixed by pipetting every 5 min to facilitate dissociation. The reaction was blocked with medium containing 10% serum and the sample was passed through a 40µm cell strainer to remove non-dissociated cells (e.g. hairs). Dissociated cells were spun down and resuspended in 1x PBS (calcium and magnesium free) with 0.04% (w/v) BSA (400 µg/ml) to a concentration of 1000 cells/µl. Following the manufacturer’s protocol and aiming for a cell recovery of 9,000 single cells per reaction, 14.9µl of each cell suspension was subjected to the single cell workflow on the 10X Genomics Chromium Controller using the Chromium Next GEM Single Cell 3ʹ Reagent Kits v3.1 (Dual Index, Revision E, with Chromium Next GEM Chip G Single Cell Kit, and Dual Index Kit TT Set A,). After GEM generation, reverse transcription and cDNA amplification, the resulting cDNA was quality controlled using a Bioanalyzer High Sensitivity DNA Assay (Agilent), and 25% of the total cDNA yield was used for fragmentation and 3’ gene expression library construction. The final libraries for Illumina short-read sequencing were analyzed using a TapeStation D1000 assay and sequenced on an Illumina NextSeq 2000 sequencer (using the NextSeq 1000/2000 P3 Reagents Kit, 100 cycles in asymmetric mode with Read1 28 bp and Read2 90 bp). After demultiplexing using the cellranger mkfastq (wrapper of bcl2fastq), 209.3 million raw reads (non-infected), 257.5 million raw reads (100 dpi) and 221.4 million raw reads (120 dpi) were assigned to the libraries.

Downstream analysis for RNAseq was performed as follows: gene abundance was quantified using salmon (v1.10.0) ^46^ with human gene annotations from gene code (version 45) for the GRCh38 genome assembly and imported using the R package tximport ^47^. Number normalization and differential expression (DE) analysis were performed using the DESeq2 package ^48^. The null variance of the Wald test statistic output from DESeq2 was re-estimated using the R package fdrtool ^49^ to calculate p-values (and adjusted using the Benjamini-Hochburg method) for the final list of differentially expressed genes. FDR (BH-adjusted p-values) ≤0.05 and log2foldchange (−1≥log2FC ≥1) were used as criteria for the final DE gene list. The R statistical computing language (version 4.1.1) was used to perform the above steps, except for gene quantification.

For spatial transcriptomic analysis on the Xenium platform (10x Genomics), 3 skin organoids for each infection condition (uninfected and infected with MCPyV for 58, 100 and 120 days) were embedded in FFPE as described above. Samples and data processing were performed using the Multimodal Segmentation Kit (nuclear staining, RNA and protein based cell staining, cell surface staining). In addition to the human multi-tissue gene panel, we measured 100 custom genes, including 8 viral targets (custom panel ID T3KHHD, see Supplementary Appendix for detailed information). Sections of 5μm thickness were placed on Xenium slides according to the manufacturer’s protocol, dried at 42°C for 3 h and placed in a desiccator at room temperature overnight, followed by deparaffinization and permeabilization to make mRNA accessible. An adjacent section was processed for RNAScope analysis to confirm the distribution of infected cells as described above. The probe hybridization mix was prepared according to the user manual (CG000582, Rev D, 10x Genomics). Xenium Nuclei Staining Buffer (10x Genomics product number: 2000762) was used to counterstain nuclei as part of the Xenium Slides & Sample Prep Reagents Kit (PN-1000460). Following the Xenium run, hematoxylin and eosin (H&E) staining was performed on the same section according to the Post-Xenium Analyzer H&E Staining User Guide (CG000613, Rev B, 10x Genomics).

### Single cell Data processing

Raw FASTQ files generated by the 10x Genomics cellranger (version 7.2.0) mkfastq pipeline were used to generate gene count matrices by the cellranger count pipeline using the human reference genome GRCh38 and GENCODE gene annotation version 32 (Ensembl 98). Custom viral transcript annotations (using “MT-” as a prefix) for both strands of the whole viral genome were added to the human genome as early and late transcripts to generate a new custom genome using cellranger mkfastq.

### Creating single-cell assay object and filtering

Cellranger output of filtered count matrices for each sample were imported into R package Seurat (v5.1. 0) ^50^ and the following filters were applied to create a scRNA-seq object for each sample using the CreateSeuratObject function: 1) genes were removed if detected in less than 10 cells; 2) cells with a total number of genes less than 200 were removed; 3) cells with a feature/gene count greater than the 99th percentile were removed; 4) cells with a proportion of mitochondrial UMIs greater than 10% were removed; and 5) cells with a total number of UMIs less than 1000 were removed.

For the fibroblast-only analysis, separate fibroblast datasets were created for each sample from the object created above, and the following sub-clustering strategies were applied in a similar manner to the original integrated scRNA-seq object with all samples.

### Normalization and integration

Normalization and variance stabilization of each scRNA-seq object was performed using the statistical modelling framework sctransform v2 ^51^ implemented in Seurat, which also removes confounding sources of variation such as: 1) total UMIs in a cell and 2) mitochondrial mapping percentage (all genes with names including “MT-”). Resulting objects for all samples were integrated stepwise using Seurat functions in order: RunPCA (with parameter npcs=30); IntegrateLayers using harmony reduction ^52^; Prior to identifying clusters of cells, a shared nearest neighbor (SNN) graph was constructed using the FindNeighbors function and used as input for clustering by the Louvain algorithm implemented in the FindClusters function. The integrated Seurat object generated from previous steps was used as input to perform non-linear dimension reduction using the Uniform Manifold Approximation and Projection (UMAP) method with the RunUMAP function.

The standard SCT assay was converted to RNA for normalization using the NormalizeData function to plot gene expression values and identify cluster markers using FindConservedMarkers. Cluster type identification and annotation was performed manually based on the expression of a set of established cell markers. Marker gene expression plots were generated using DotPlot and FeaturePlot functions and ggplot2^53^ using R version 4.4.1 (https://www.R-project.org/). Gene expression immunity score was calculated using the AddModuleScore function using only selected genes from the bulk RNA-seq analysis.

### Spatial transcriptomic data processing

Spatial transcriptomic data were analyzed in R using the VoltRon package ^54^ and packages from tidyverse ^55^, DESeq2 ^48^ and apeglm ^56^. MCPyV early and late gene expression was determined using a combination of 4 probes detecting early genes (MCPyV_Early_2, MCPyV_Early_canonical_1, MCPyV_Early_canonical_2 and MCPyV_sT) and a probe specific for the MCPyV miR-M1 (MCPyV_miR2L). A cell was defined as positive for early viral gene expression if at least 3 of the 4 early gene probes were detected simultaneously with at least one count each. A cell was defined as positive for late MCPyV gene expression when 3 counts of MCPyV_miR2L were detected simultaneously with the early genes. Cells with 10 or fewer cellular RNA counts were filtered out. The code is available on github.

### Statistical analysis

Statistical analysis was performed using GraphPad Prism (version 10, GraphPad Software, San Diego,CA).

---

**Supplementary Figure S1 Characterization of SkO differentiation**. **a,** Schematic representation of the SkO generation protocol with timeline (created with BioRender.com); **b,** Brightfield images of SkOs at the indicated time points of differentiation. **c,** Whole mount immunostaining of SkOs at the indicated time points of differentiation. Top: KRT17: epidermis and outer root sheath of organoids (magenta); KRT15: periderm and basal layers of the skin (yellow); PDGFRα: dermal fibroblasts (green). Middle: KI67: proliferating cells (magenta); TUJI: neuronal cells and Schwann cells (yellow); LHX2: hair placodes and bulge region (cyan). Bottom: SOX2: dermal condensates, dermal papilla, melanocytes and Merkel cells (cyan); ECAD: epithelium (magenta); KRT20: Merkel cells (yellow). Hoechst counterstain for nuclei (blue). Where indicated, images show the maximum intensity projection (MIP) of multiple z-stack acquisitions. Elsewhere a single z-stack is shown. Dashed boxes indicate the magnified regions at the bottom. Brightness and contrast have been adjusted independently for each image to optimize visibility. Scale bar: 500 μm in the 15 to 120dod SkOs, 100 μm in the 0dod and 3dod and zoom-ins.

**Supplementary Figure S2 MCPyV infection at early and late stages of SkO differentiation. a,** Schematic representation of the time course of infection in SkOs infected at 146dod. **b,** Whole-mount immunostaining of end-stage differentiated SkOs infected with MCPyV at the indicated time points. **c,** Schematic representation of the time course of infection in SkOs infected at 45dod. **d,** Whole mount immunostaining of early-stage differentiated SkOs infected with MCPyV at the indicated time points. In b and d, MCPyV VP1 stains for the virus capsid protein (white) and Hoechst counterstains nuclei (blue). Where indicated, the images show the maximum intensity projection (MIP) of multiple z-stack acquisitions. Elsewhere a single z-stack is shown. Yellow dashed boxes indicate the magnified regions shown below. The dashed line in the single z-stack magnification indicates the boundary between the dermis and the SkOs hair follicles, stratified epithelium, and tail. Scale bar: 500 μm in the main images, 200 μm in the magnifications.

**Supplementary Figure S3 Time course of MCPyV infection in SkOs. a,** Representation of the time course experiment; SkOs were infected with 1.5×10E08 genome equivalents per SkO at 30dod and harvested at subsequent time points (blue shading), or infected at different time points along the differentiation and harvested at the end point (150dod, green shading). **b,** Whole-mount immunostaining of SkOs infected with MCPyV for 4 to 120 days. VP1 (green) stains the MCPyV capsid protein, KRT17 (magenta), stains the epidermis; Hoechst (blue) counterstains nuclei. On the outer left and right sides of the images, the day of differentiation at harvest/infection is indicated for each sample. Where indicated, the images show the maximum intensity projection (MIP) of multiple z-stack acquisitions. Elsewhere a single z-stack is shown. The dotted white insets indicate the location of the magnifications in columns one and four. Scale bar 500 μm in the main images, 100 μm in the magnifications.

**Supplementary Figure S4 Differences in infection in early and late differentiated SkOs. a,** Schematic representation of the time course of infection in SkOs infected at early vs late stage of differentiation, as in Figure S3. **b,** SkOs were formalin-fixed paraffin-embedded (FFPE) and serial sections were labelled by in situ hybridization DNAScope (red) visualizing the MCPyV genome (first column), immunohistochemistry for MCPyV LT (Ab3) antigen or PDGFRα (dermal fibroblasts, second and fourth columns, respectively), and immunofluorescence for ECAD (epithelium, magenta), MCPyV VP1 (green), DAPI (nuclei, blue, third column). Dashed boxes indicate the magnified regions. Scale bar: 200 μm.

**Supplementary Figure S5 Productive infection of MCPyV in SkOs. a,** Electron microscopy (EM) of negative stained MCPyV virions produced in 293 Lentix cells (293-MCPyV, left) or isolated from the SkOs supernatant (SkO-MCPyV, right). **b,** Immunostaining of nHDF infected for five days with 2×10E05 genome equivalents of 293-MCPyV or SkO-MCPyV per cell. VP1 stains for MCPyV capsid protein (green), LT CM2B4 stains for MCPyV LT (magenta); DAPI counterstains nuclei. Scale bar: 200 μm in the main images, 50 μm zoom-in. **c,** Whole-mount immunostaining of SkOs infected with 293-MCPyV (top) or SKO-MCPyV (bottom) for the indicated time points and harvested at day 150 of differentiation. Scale bars: 500 μm.

**Supplementary Figure S6 Electron microscopy of MCPyV infected organoids.** Transmission electron microscopy (TEM) images of a 150dod/100dpi SkO. **a,** Representative images showing the location of infected cells in the SkO. Dashed insets highlight the area magnified in the subsequent images. HF: hair follicles; LC: lipid cells. **b,** 40-60 nm electron-dense virus particles in cytoplasmic vesicles. **c and d,** lytic cells showing virus particles in the nucleus. Dashed insets in d, indicate the magnified region to the right. Red arrowhead points to tubular structures associated with MCPyV virions. **e,** cell debris containing organized viral particles.

**Supplementary Figure S7 Schematic representation of the sequencing analysis on MCPyV-infected SkOs. a,** Schematic representation of the infection time course in SkOs. **b,** Samples used for bulk RNA-seq analysis. mRNA was isolated from four SkOs per condition and an equal amount of mRNA from two SkOs was pooled. **c,** For scRNA-seq analysis single cell dissociation was performed on four SkOs per condition and pooled together. 9,000 cells from each pool were further processed for library preparation. **d,** For spatial transcriptomics three SkOs per condition were embedded in FFPE blocks and further processed for spatial analysis.

**Supplementary Figure S8 scRNA-seq cluster definition.** For each cluster, the gene expression of three well-established cell type markers was projected onto the UMAP plot to identify the different cell type populations. The color code refers to the level of gene expression, with red indicating maximum gene expression and blue indicating low or no expression.

**Supplementary Figure S9 Characterization of SkOs by scRNA-seq and spatial transcriptomics. a,** UMAP plots of 150day-old SkOs from scRNA-seq (left) and spatial transcriptomic (right) datasets, showing the different clusters identified. **b,** Spatial plot of a non-infected SkO; colors represent the different cell types. **c,** Spatial plot showing the location of papillary fibroblasts (white).

**Supplementary Figure S10 MCPyV infected cells in the SkOs.** Early (top) and late (left) MCPyV gene expression in different cell types. For each cluster, the total number of cells/number of infected cells is shown. The percentage of infected cells is shown in parentheses.

**Supplementary Figure S11 MCPyV infection in SkOs activates the IFNβ pathway. a,** Principal component analysis of non-infected SkOs (red) and SkOs infected for 100 (light blue) and 120 days (blue) from the bulk RNA-seq dataset. **b,** Volcano plot depicting all the deregulated genes (DEGs) in SkOs infected for 100 (left) or 120 days (right) compared to non-infected SkOs. Genes that are significantly (padj. ≤0.05) up-(log2FC ≥1) or down-regulated (log2FC ≤-1) are shown as red or blue circles, respectively. **c,** Feature plot showing normalized intensity level of the indicated ISGs in non-infected and MCPyV-infected SkOs. The color code refers to the intensity of gene expression, with blue indicating maximum gene expression and white indicating low or no expression.

**Supplementary Figure S12** Whole-mount immunostaining of 170dod/60dpi and treated with vehicle control (DMSO), IFNβ or ruxolitinib (RUX), throughout the infection. KRT17 staining for epidermis and outer root sheath of the hairs (magenta), MCPyV VP1 staining for the virus capsid protein (green), and Hoechst counterstaining for nuclei (blue). The images show the maximum intensity projection (MIP) of multiple z-stack acquisitions. Dashed boxes indicate the magnified regions. Scale bar: 500 μm in the main images, 100 μm in the zoom-ins.

## Acknowledgements

We thank Robert Shelansky, Aishwarya Konnur, Jana Lalakova, Morgane Rouault (10xGenomics) for performing the primary spatial experiment in this manuscript and for their technical support. We are grateful to Vera Wolf and Kathrein Permien (Charité, Berlin) for their valuable technical assistance. We also thank the UKE imaging facility (umif, UKE) and Christian Conze (LIV, light microscopy and image analysis unit) for their excellent support. We acknowledge Claudia Schmidt, Nina Bracker, and Josephine Bonath (UKE) for their assistance with iPSC and organoid culture. We are grateful to Chris Buck (NIH, USA) and James DeCaprio (Dana Farber Institute, USA) for generously providing antibodies. This project was funded by the Deutsche Forschungsgemeinschaft (DFG, German Research Foundation) in the framework of the Research Unit FOR5200 DEEP-DV (443644894) projects FI 782/ 7-1 (N.F.), GR 3318/5-1 (A.G.).

## Contributions

S.A., M.C.-S., E.W., M.L., A.G. and N.F. conceptualized the study. S.A., M.C.-S., E.W., T.G., P.B., developed the methodology. S.A., E.W., L.R, V.B., L.A., C.S., R.R., S.K. conducted the investigations, S.A., E.W., S.V., T.G., conducted the formal analysis. M.D., A. H. provided reagents and techniques. S.A. and N.F. wrote the original draft. A.G., M.L. and N.F. acquired funding.

## References

1. Becker, J.C., et al. Epidemiology, biology and therapy of Merkel cell carcinoma: conclusions from the EU project IMMOMEC. Cancer Immunol Immunother 67, 341–351 (2018).

2. DeCaprio, J.A. Molecular Pathogenesis of Merkel Cell Carcinoma. Annu Rev Pathol 16, 69–91 (2021).

3. Feng, H., Shuda, M., Chang, Y. & Moore, P.S. Clonal integration of a polyomavirus in human Merkel cell carcinoma. Science 319, 1096–1100 (2008).

4. Houben, R., et al. Merkel cell polyomavirus-infected Merkel cell carcinoma cells require expression of viral T antigens. J Virol 84, 7064–7072 (2010).

5. Shuda, M., et al. T antigen mutations are a human tumor-specific signature for Merkel cell polyomavirus. Proc Natl Acad Sci U S A 105, 16272–16277 (2008).

6. Goh, G., et al. Mutational landscape of MCPyV-positive and MCPyV-negative Merkel cell carcinomas with implications for immunotherapy. Oncotarget 7, 3403–3415 (2016).

7. Harms, P.W., et al. The Distinctive Mutational Spectra of Polyomavirus-Negative Merkel Cell Carcinoma. Cancer Res 75, 3720–3727 (2015).

8. Starrett, G.J., et al. Clinical and molecular characterization of virus-positive and virus-negative Merkel cell carcinoma. Genome Med 12, 30 (2020).

9. Grundhoff, A. & Fischer, N. Merkel cell polyomavirus, a highly prevalent virus with tumorigenic potential. Curr Opin Virol 14, 129–137 (2015).

10. Feng, H., et al. Cellular and viral factors regulating Merkel cell polyomavirus replication. PLoS One 6, e22468 (2011).

11. Liu, W., et al. Identifying the Target Cells and Mechanisms of Merkel Cell Polyomavirus Infection. Cell Host Microbe 19, 775–787 (2016).

12. Neumann, F., et al. Replication, gene expression and particle production by a consensus Merkel Cell Polyomavirus (MCPyV) genome. PLoS One 6, e29112 (2011).

13. Ohnezeit, D., et al. Merkel cell polyomavirus small tumor antigen contributes to immune evasion by interfering with type I interferon signaling. PLoS Pathog 20, e1012426 (2024).

14. Liu, W., Krump, N.A., Buck, C.B. & You, J. Merkel Cell Polyomavirus Infection and Detection. J Vis Exp (2019).

15. Spurgeon, M.E., et al. The Merkel Cell Polyomavirus T Antigens Function as Tumor Promoters in Murine Skin. Cancers (Basel) 13(2021).

16. Verhaegen, M.E., et al. Direct cellular reprogramming enables development of viral T antigen-driven Merkel cell carcinoma in mice. J Clin Invest 132(2022).

17. Verhaegen, M.E., et al. Merkel Cell Polyomavirus Small T Antigen Initiates Merkel Cell Carcinoma-like Tumor Development in Mice. Cancer Res 77, 3151–3157 (2017).

18. Li, P., et al. Mpox virus infection and drug treatment modelled in human skin organoids. Nat Microbiol 8, 2067–2079 (2023).

19. Ma, J., et al. Establishment of Human Pluripotent Stem Cell-Derived Skin Organoids Enabled Pathophysiological Model of SARS-CoV-2 Infection. Adv Sci (Weinh) 9, e2104192 (2022).

20. Lee, J., et al. Hair-bearing human skin generated entirely from pluripotent stem cells. Nature 582, 399–404 (2020).

21. Lee, J., et al. Generation and characterization of hair-bearing skin organoids from human pluripotent stem cells. Nat Protoc 17, 1266–1305 (2022).

22. Theiss, J.M., et al. A Comprehensive Analysis of Replicating Merkel Cell Polyomavirus Genomes Delineates the Viral Transcription Program and Suggests a Role for mcv-miR-M1 in Episomal Persistence. PLoS Pathog 11, e1004974 (2015).

23. Jung, S.Y., et al. Wnt-activating human skin organoid model of atopic dermatitis induced by Staphylococcus aureus and its protective effects by Cutibacterium acnes. iScience 25, 105150 (2022).

24. Ramovs, V., et al. Characterization of the epidermal-dermal junction in hiPSC-derived skin organoids. Stem Cell Reports 17, 1279–1288 (2022).

25. Becker, M., et al. Infectious Entry of Merkel Cell Polyomavirus. J Virol 93(2019).

26. Dupzyk, A. & Tsai, B. Bag2 Is a Component of a Cytosolic Extraction Machinery That Promotes Membrane Penetration of a Nonenveloped Virus. J Virol 92(2018).

27. Dupzyk, A., Williams, J.M., Bagchi, P., Inoue, T. & Tsai, B. SGTA-Dependent Regulation of Hsc70 Promotes Cytosol Entry of Simian Virus 40 from the Endoplasmic Reticulum. J Virol 91(2017).

28. Nakanishi, A., Clever, J., Yamada, M., Li, P.P. & Kasamatsu, H. Association with capsid proteins promotes nuclear targeting of simian virus 40 DNA. Proc Natl Acad Sci U S A 93, 96–100 (1996).

29. Erickson, K.D., et al. Virion assembly factories in the nucleus of polyomavirus-infected cells. PLoS Pathog 8, e1002630 (2012).

30. Philippeos, C., et al. Spatial and Single-Cell Transcriptional Profiling Identifies Functionally Distinct Human Dermal Fibroblast Subpopulations. J Invest Dermatol 138, 811–825 (2018).

31. Sole-Boldo, L., et al. Single-cell transcriptomes of the human skin reveal age-related loss of fibroblast priming. Commun Biol 3, 188 (2020).

32. Tabib, T., Morse, C., Wang, T., Chen, W. & Lafyatis, R. SFRP2/DPP4 and FMO1/LSP1 Define Major Fibroblast Populations in Human Skin. J Invest Dermatol 138, 802–810 (2018).

33. Ahlers, J.M.D., et al. Single-Cell RNA Profiling of Human Skin Reveals Age-Related Loss of Dermal Sheath Cells and Their Contribution to a Juvenile Phenotype. Front Genet 12, 797747 (2021).

34. Wang, R., Yang, J.F., Senay, T.E., Liu, W. & You, J. Characterization of the Impact of Merkel Cell Polyomavirus-Induced Interferon Signaling on Viral Infection. J Virol 97, e0190722 (2023).

35. Spurgeon, M.E., et al. Merkel cell polyomavirus large T antigen binding to pRb promotes skin hyperplasia and tumor development. PLoS Pathog 18, e1010551 (2022).

36. Verhaegen, M.E., et al. Merkel cell polyomavirus small T antigen is oncogenic in transgenic mice. J Invest Dermatol 135, 1415–1424 (2015).

37. Foulongne, V., et al. Merkel cell polyomavirus in cutaneous swabs. Emerg Infect Dis 16, 685–687 (2010).

38. Wieland, U., Mauch, C., Kreuter, A., Krieg, T. & Pfister, H. Merkel cell polyomavirus DNA in persons without merkel cell carcinoma. Emerg Infect Dis 15, 1496–1498 (2009).

39. Sahragard, I., et al. Impact of BK Polyomavirus NCCR variations in post kidney transplant outcomes. Gene 913, 148376 (2024).

40. Carter, J.J., et al. Identification of an overprinting gene in Merkel cell polyomavirus provides evolutionary insight into the birth of viral genes. Proc Natl Acad Sci U S A 110, 12744–12749 (2013).

41. Salisbury, N.J.H., et al. Polyomavirus ALTOs, but not MTs, downregulate viral early gene expression by activating the NF-kappaB pathway. Proc Natl Acad Sci U S A 121, e2403133121 (2024).

42. Wang, R., et al. Merkel cell polyomavirus protein ALTO modulates TBK1 activity to support persistent infection. PLoS Pathog 20, e1012170 (2024).

43. Yang, R., et al. Characterization of ALTO-encoding circular RNAs expressed by Merkel cell polyomavirus and trichodysplasia spinulosa polyomavirus. PLoS Pathog 17, e1009582 (2021).

44. Rahmani, W., Sinha, S. & Biernaskie, J. Immune modulation of hair follicle regeneration. NPJ Regen Med 5, 9 (2020).

45. Schowalter, R.M., Pastrana, D.V., Pumphrey, K.A., Moyer, A.L. & Buck, C.B. Merkel cell polyomavirus and two previously unknown polyomaviruses are chronically shed from human skin. Cell Host Microbe 7, 509–515 (2010).

46. Patro, R., Duggal, G., Love, M.I., Irizarry, R.A. & Kingsford, C. Salmon provides fast and bias-aware quantification of transcript expression. Nat Methods 14, 417–419 (2017).

47. Soneson, C., Love, M.I. & Robinson, M.D. Differential analyses for RNA-seq: transcript-level estimates improve gene-level inferences. F1000Res 4, 1521 (2015).

48. Love, M.I., Huber, W. & Anders, S. Moderated estimation of fold change and dispersion for RNA-seq data with DESeq2. Genome Biol 15, 550 (2014).

49. Strimmer, K. fdrtool: a versatile R package for estimating local and tail area-based false discovery rates. Bioinformatics 24, 1461–1462 (2008).

50. Hao, Y., et al. Dictionary learning for integrative, multimodal and scalable single-cell analysis. Nat Biotechnol 42, 293–304 (2024).

51. Choudhary, S. & Satija, R. Comparison and evaluation of statistical error models for scRNA-seq. Genome Biol 23, 27 (2022).

52. Korsunsky, I., et al. Fast, sensitive and accurate integration of single-cell data with Harmony. Nat Methods 16, 1289–1296 (2019).

53. 53. Wickham, H. Ggplot2. Springer International Publishing (2016).

54. Manukyan, A., et al. VoltRon: A Spatial Omics Analysis Platform for Multi-Resolution and Multiomics Integration using Image Registration. bioRxiv, 2023.2012.2015.571667 (2023).

55. Wickham, H., et al. Welcome to the Tidyverse. Journal of Open Source Software (2019).

56. Zhu, A., Ibrahim, J.G. & Love, M.I. Heavy-tailed prior distributions for sequence count data: removing the noise and preserving large differences. Bioinformatics 35, 2084–2092 (2019).

