## Supplementary Fig S1-S12 for "Merkel cell polyomavirus infection and persistence modelled in skin organoids"

Figure_S1

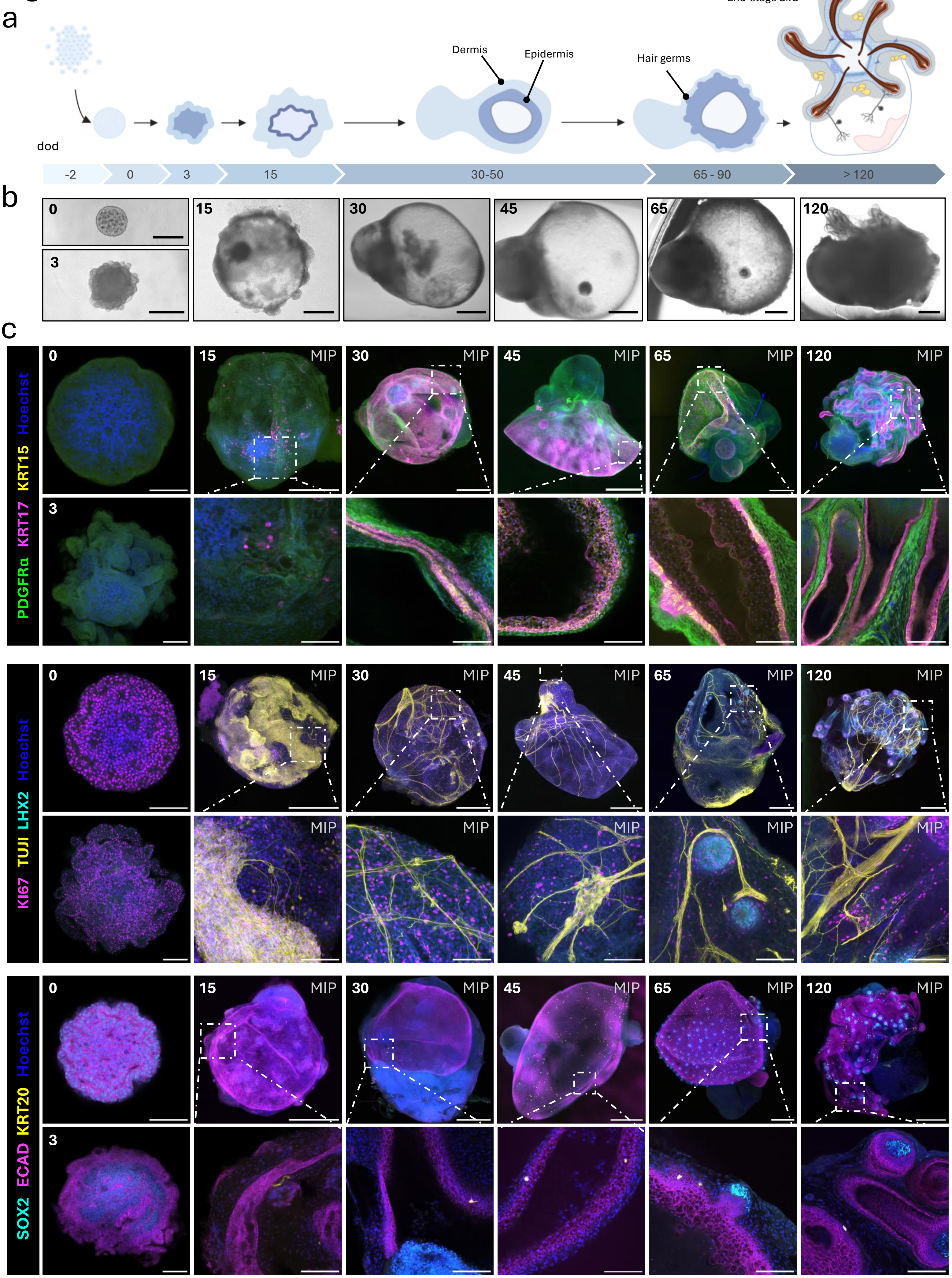
a
End-stage SkO
Dermis
Epidermis
Hair germs
dod
-2
0
3
15
30-50
65 - 90
> 120
b
0
3
15
30
45
65
120
c
PDGFRα KRT17 KRT15 Hoechst
KI67 TUJI LHX2 Hoechst
SOX2 ECAD KRT20 Hoechst
MIP

Figure_S2

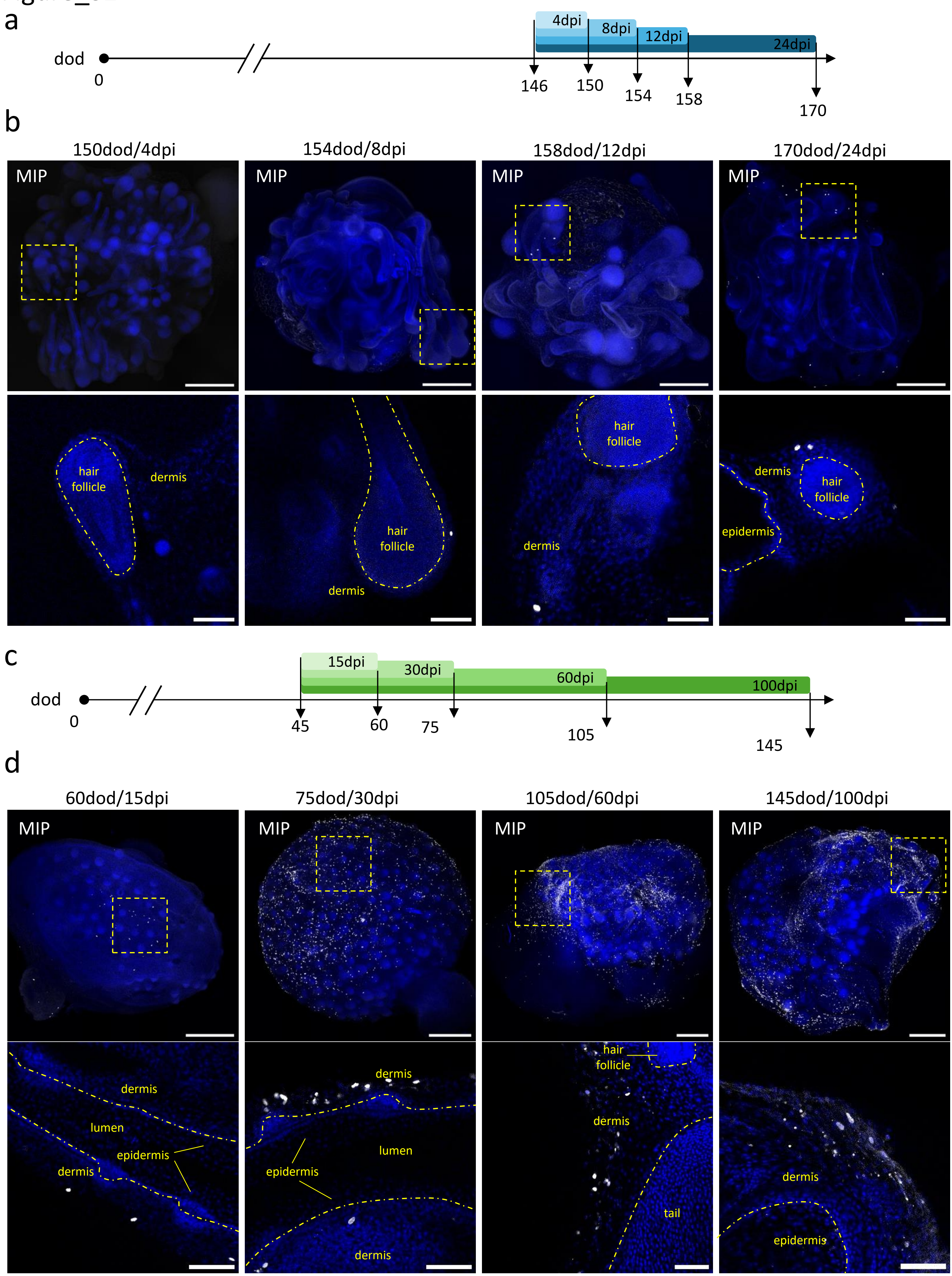
a
4dpi
8dpi
12dpi
24dpi
dod
0
146
150
154
158
170
b
150dod/4dpi
154dod/8dpi
158dod/12dpi
170dod/24dpi
MIP
hair follicle
dermis
epidermis
c
15dpi
30dpi
60dpi
100dpi
dod
0
45
60
75
105
145
d
60dod/15dpi
75dod/30dpi
105dod/60dpi
145dod/100dpi
MIP
dermis
lumen
epidermis
hair follicle
tail

Figure_S3

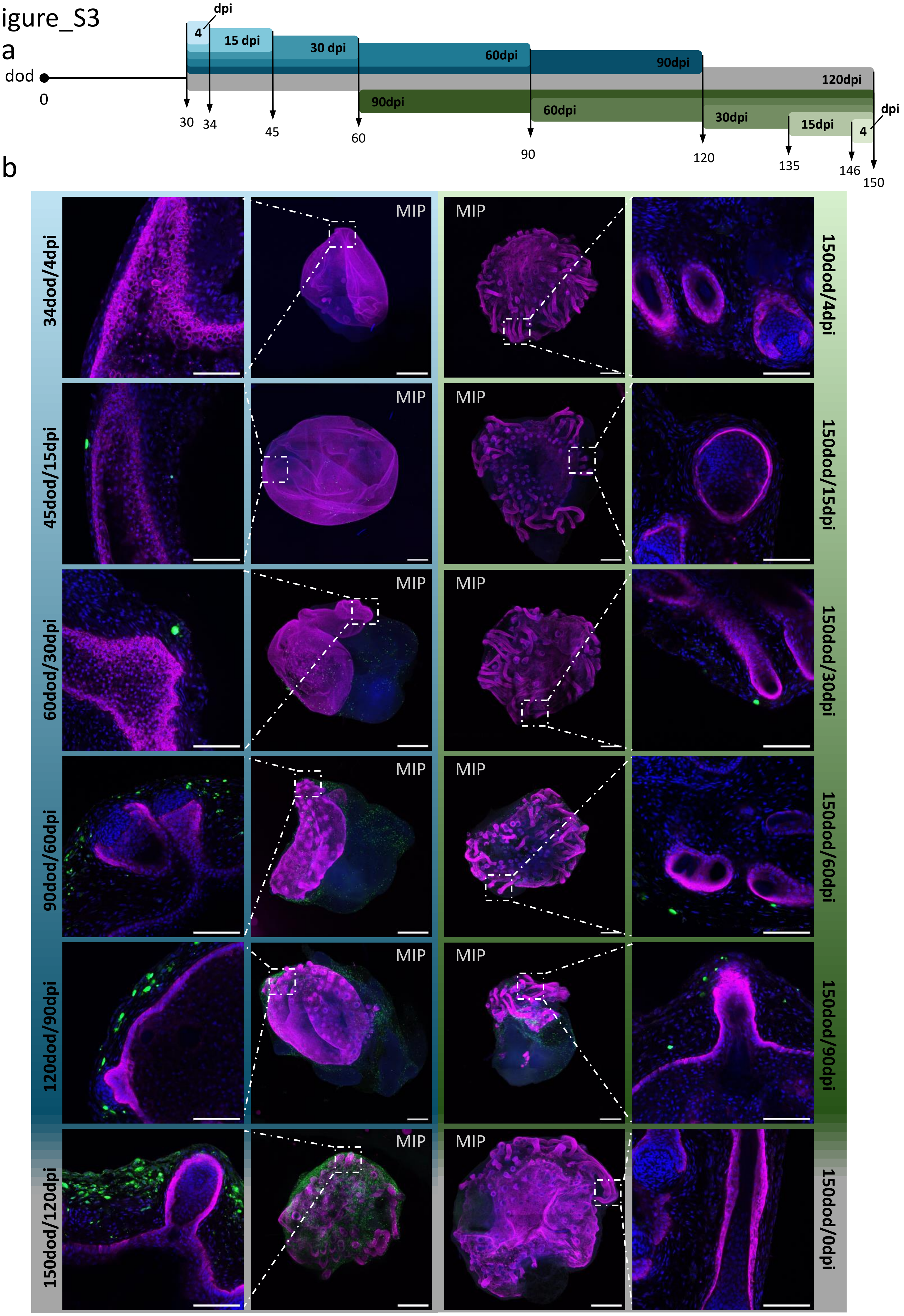

a
dod
0
dpi
4
15 dpi
30 dpi
60dpi
90dpi
120dpi
90dpi
60dpi
30dpi
15dpi
4
dpi
30
34
45
60
90
120
135
146
150
b
34dod/4dpi
45dod/15dpi
60dod/30dpi
90dod/60dpi
120dod/90dpi
150dod/120dpi
MIP
150dod/4dpi
150dod/15dpi
150dod/30dpi
150dod/60dpi
150dod/90dpi
150dod/0dpi

### Figure_S4

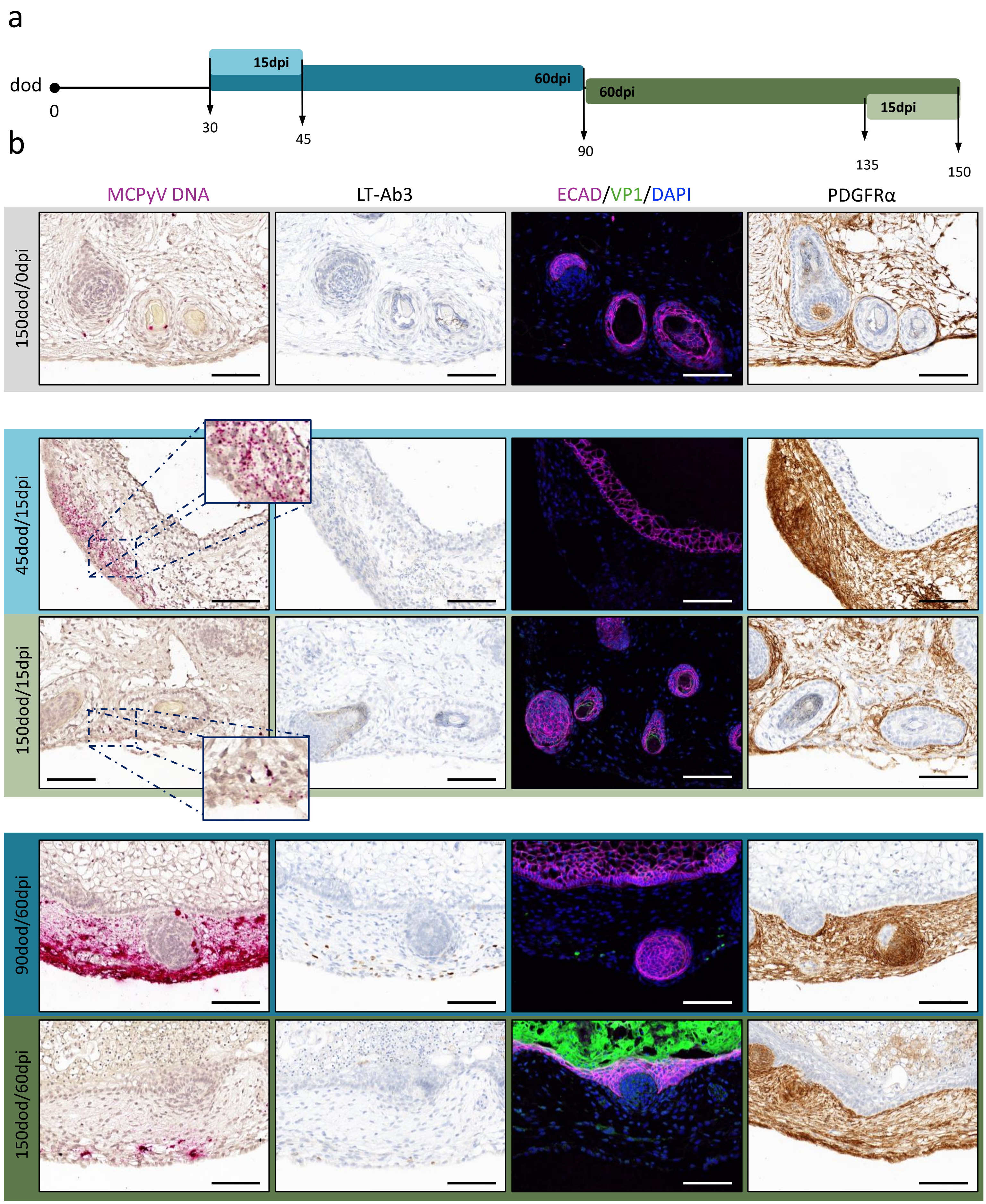
a
dod
0
30
45
90
135
150
15dpi
60dpi
60dpi
15dpi
b
MCPyV DNA
LT-Ab3
ECAD/VP1/DAPI
PDGFRα
150dod/0dpi
45dod/15dpi
150dod/15dpi
90dod/60dpi
150dod/60dpi

Figure_S5

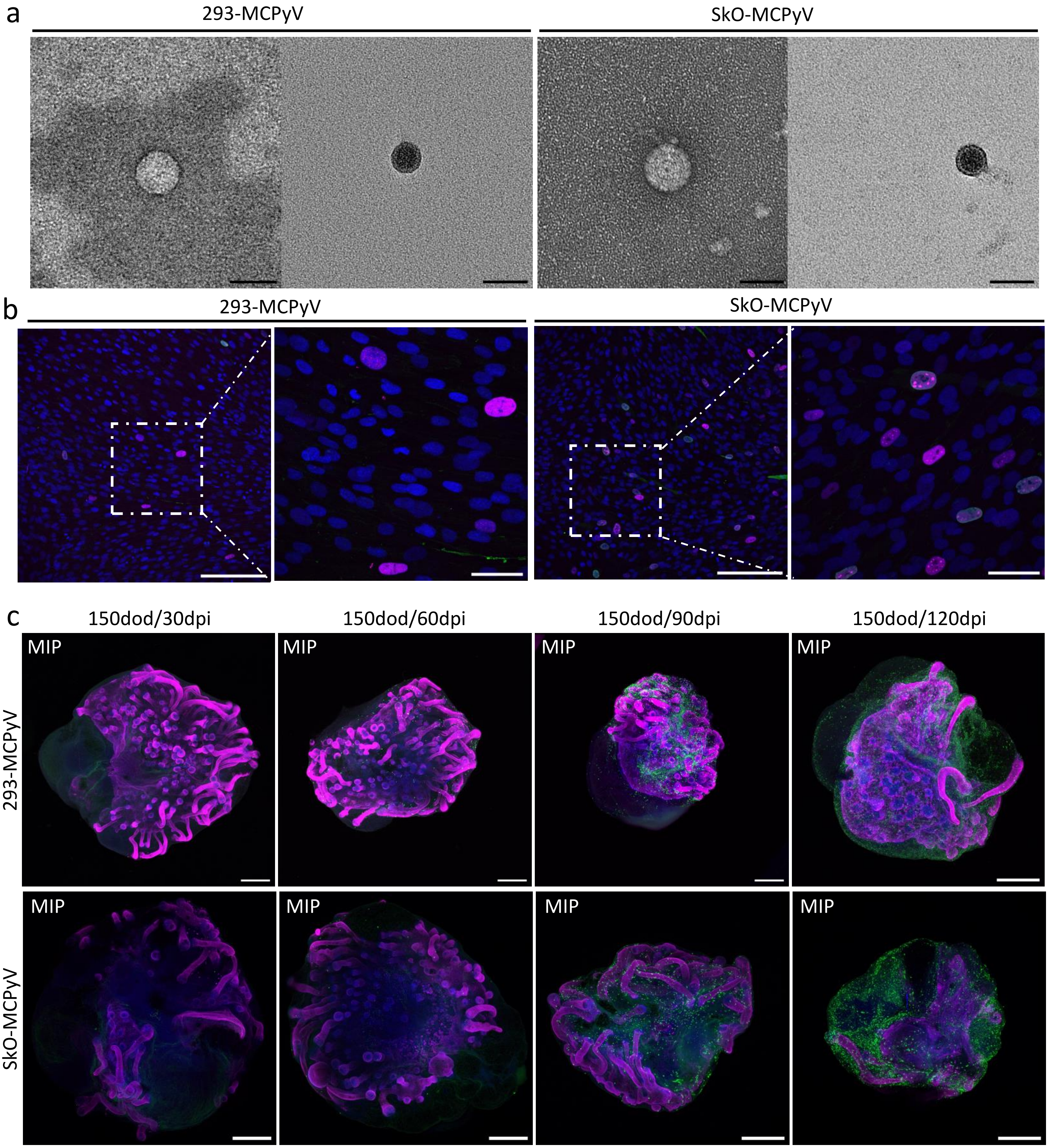
a
293-MCPyV
SkO-MCPyV
b
293-MCPyV
SkO-MCPyV
c
150dod/30dpi
150dod/60dpi
150dod/90dpi
150dod/120dpi
293-MCPyV
SkO-MCPyV
MIP

Figure_S6

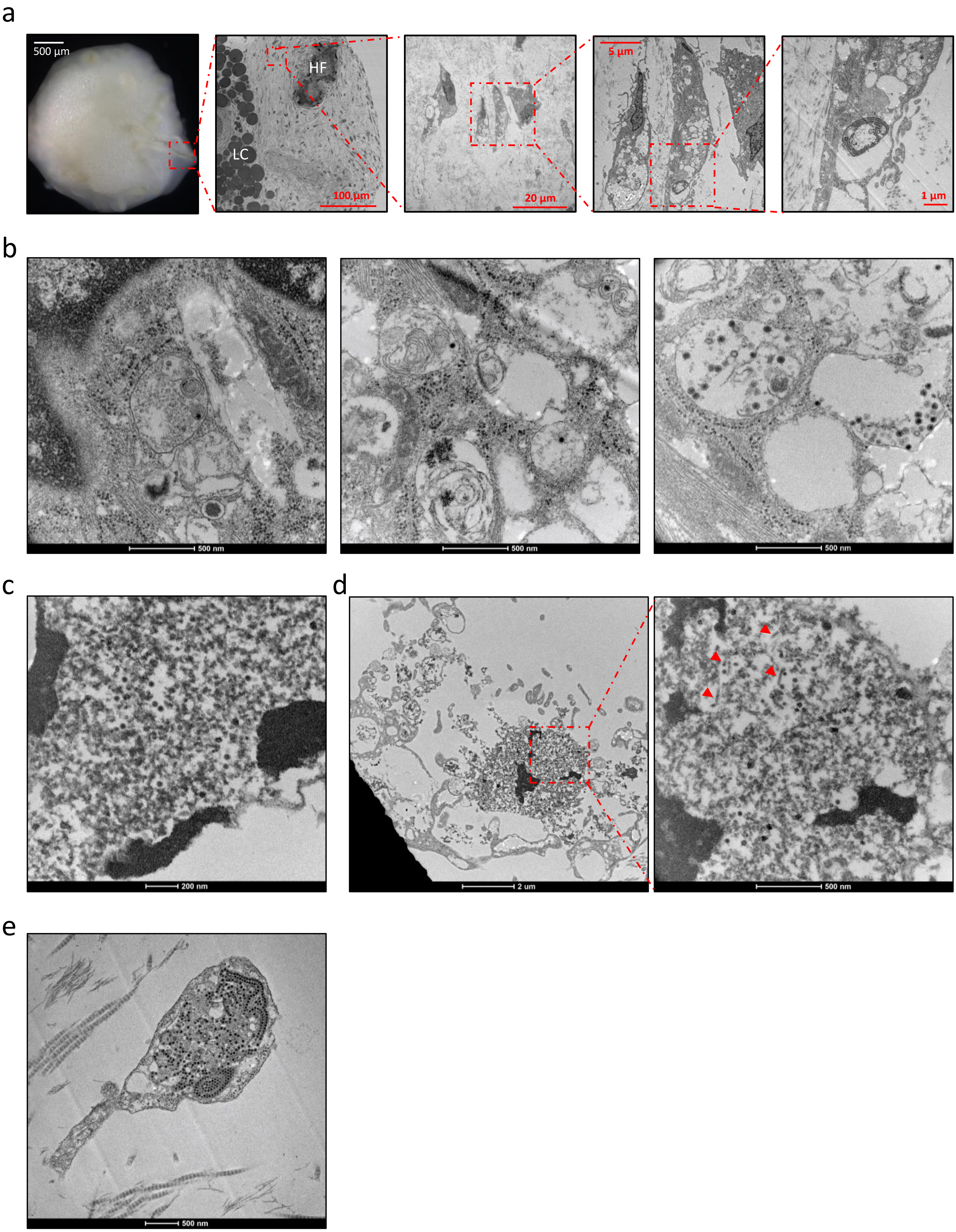
a
500 µm
HF
LC
100 µm
20 µm
5 µm
1 µm
b
500 nm
500 nm
500 nm
c
200 nm
d
2 um
500 nm
e
500 nm

Figure_S7

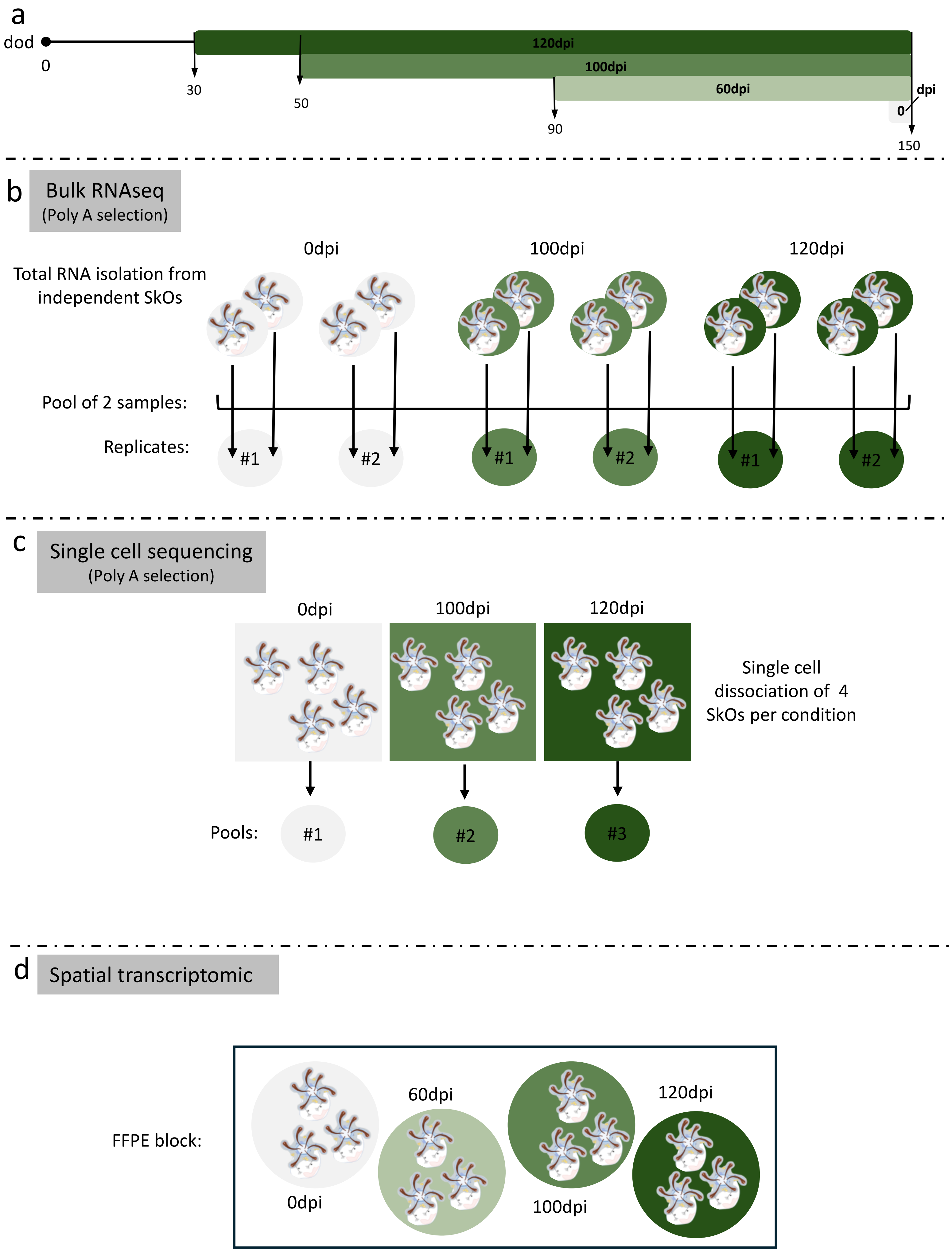
a
dod
0
30
50
90
150
120dpi
100dpi
60dpi
dpi
0
b
Bulk RNAseq
(Poly A selection)
0dpi
100dpi
120dpi
Total RNA isolation from independent SkOs
Pool of 2 samples:
Replicates:
#1
#2
#1
#2
#1
#2
c
Single cell sequencing
(Poly A selection)
0dpi
100dpi
120dpi
Single cell dissociation of 4 SkOs per condition
Pools:
#1
#2
#3
d
Spatial transcriptomic
FFPE block:
60dpi
120dpi
0dpi
100dpi

Figure_S8

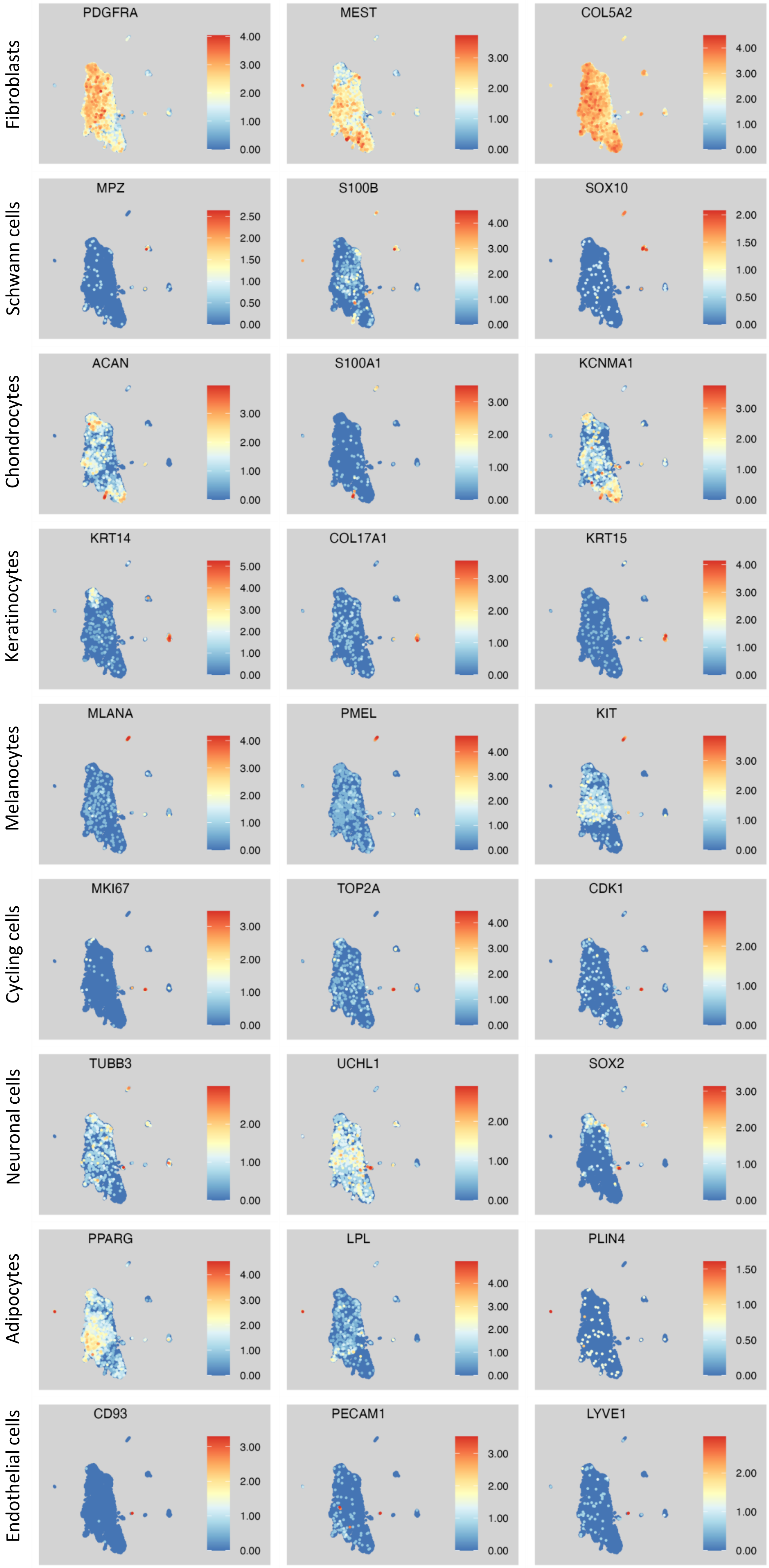

Fibroblasts
PDGFRA
MEST
COL5A2
Schwann cells
MPZ
S100B
SOX10
Chondrocytes
ACAN
S100A1
KCNMA1
Keratinocytes
KRT14
COL17A1
KRT15
Melanocytes
MLANA
PMEL
KIT
Cycling cells
MKI67
TOP2A
CDK1
Neuronal cells
TUBB3
UCHL1
SOX2
Adipocytes
PPARG
LPL
PLIN4
Endothelial cells
CD93
PECAM1
LYVE1

Figure_S9

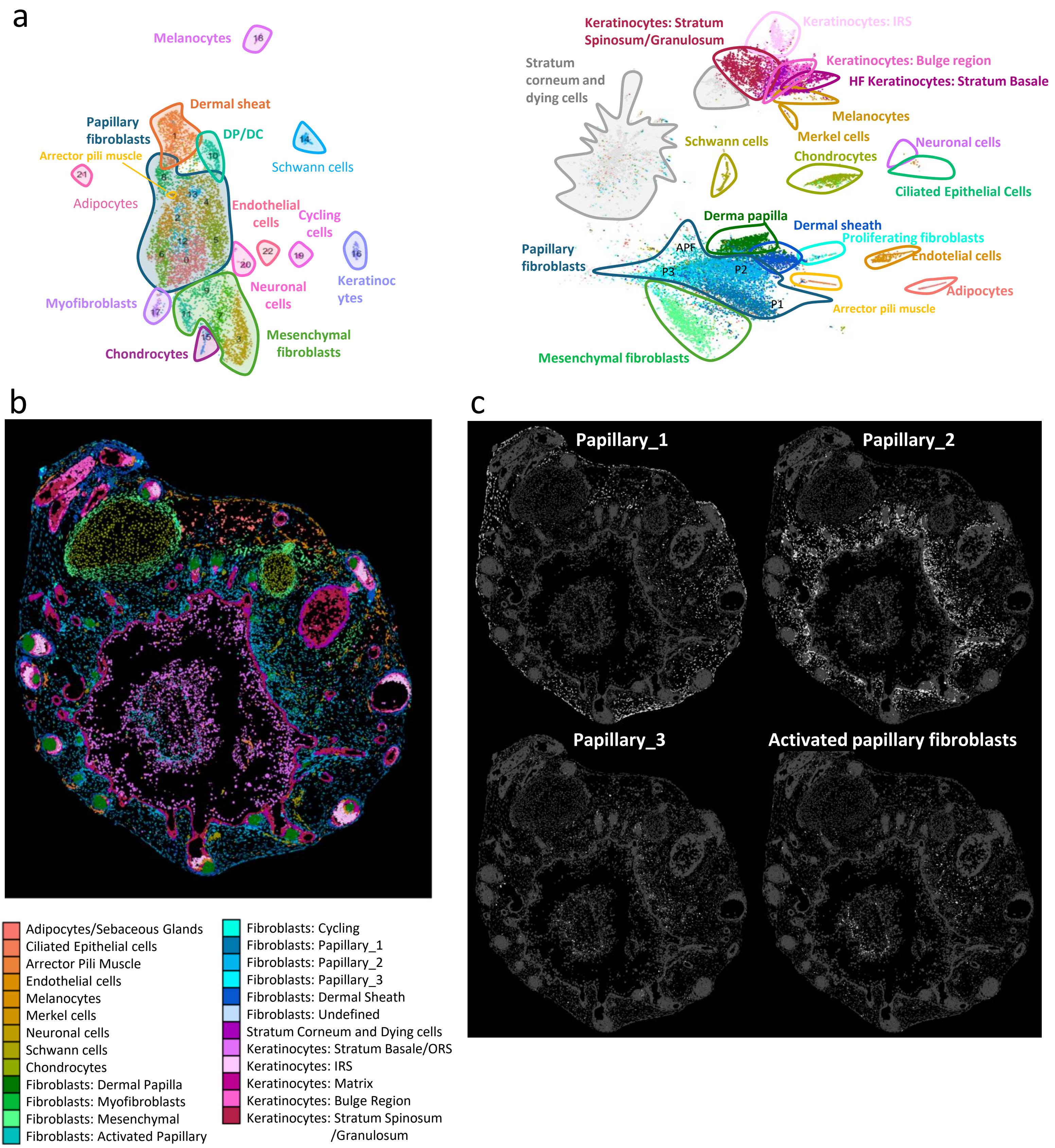
a
Melanocytes
Dermal sheat
Papillary fibroblasts
DP/DC
Arrector pili muscle
Schwann cells
Adipocytes
Endothelial cells
Cycling cells
Keratinocytes
Myofibroblasts
Neuronal cells
Mesenchymal fibroblasts
Chondrocytes
Keratinocytes: Stratum Spinosum/Granulosum
Keratinocytes: IRS
Keratinocytes: Bulge region
HF Keratinocytes: Stratum Basale
Stratum corneum and dying cells
Melanocytes
Merkel cells
Schwann cells
Chondrocytes
Neuronal cells
Ciliated Epithelial Cells
Derma papilla
Dermal sheath
Papillary fibroblasts
APF
P3
P2
P1
Proliferating fibroblasts
Endotelial cells
Adipocytes
Arrector pili muscle
Mesenchymal fibroblasts
b
Adipocytes/Sebaceous Glands
Ciliated Epithelial cells
Arrector Pili Muscle
Endothelial cells
Melanocytes
Merkel cells
Neuronal cells
Schwann cells
Chondrocytes
Fibroblasts: Dermal Papilla
Fibroblasts: Myofibroblasts
Fibroblasts: Mesenchymal
Fibroblasts: Activated Papillary
Fibroblasts: Cycling
Fibroblasts: Papillary_1
Fibroblasts: Papillary_2
Fibroblasts: Papillary_3
Fibroblasts: Dermal Sheath
Fibroblasts: Undefined
Stratum Corneum and Dying cells
Keratinocytes: Stratum Basale/ORS
Keratinocytes: IRS
Keratinocytes: Matrix
Keratinocytes: Bulge Region
Keratinocytes: Stratum Spinosum /Granulosum
c
Papillary_1
Papillary_2
Papillary_3
Activated papillary fibroblasts

### Figure_S10

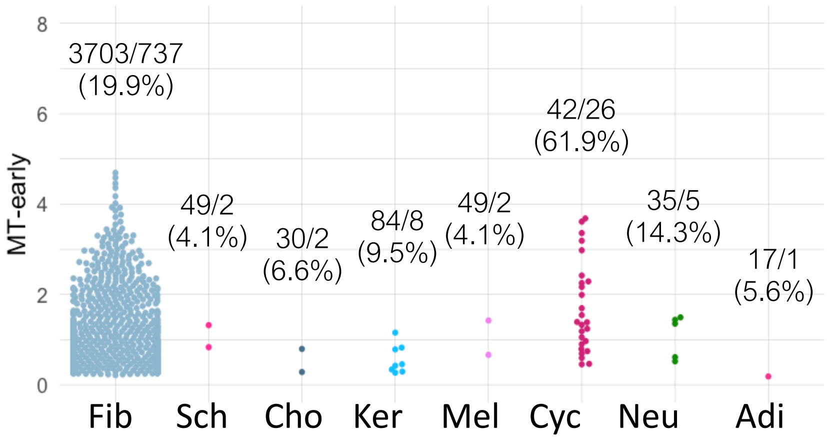

MT-early
8
6
4
2
0
3703/737
(19.9%)
49/2
(4.1%)
30/2
(6.6%)
84/8
(9.5%)
49/2
(4.1%)
42/26
(61.9%)
35/5
(14.3%)
17/1
(5.6%)
Fib
Sch
Cho
Ker
Mel
Cyc
Neu
Adi


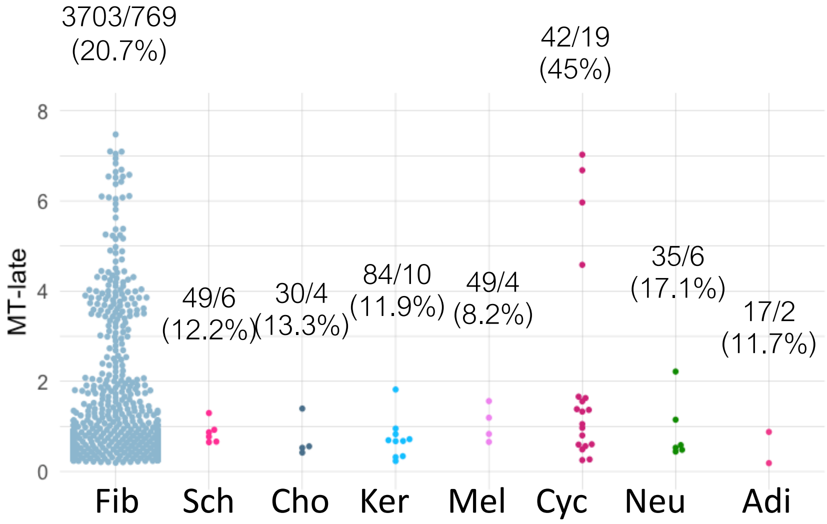

3703/769
(20.7%)
42/19
(45%)
MT-late
8
6
4
2
0
49/6
(12.2%)
30/4
(13.3%)
84/10
(11.9%)
49/4
(8.2%)
35/6
(17.1%)
17/2
(11.7%)
Fib
Sch
Cho
Ker
Mel
Cyc
Neu
Adi

### Figure_S11

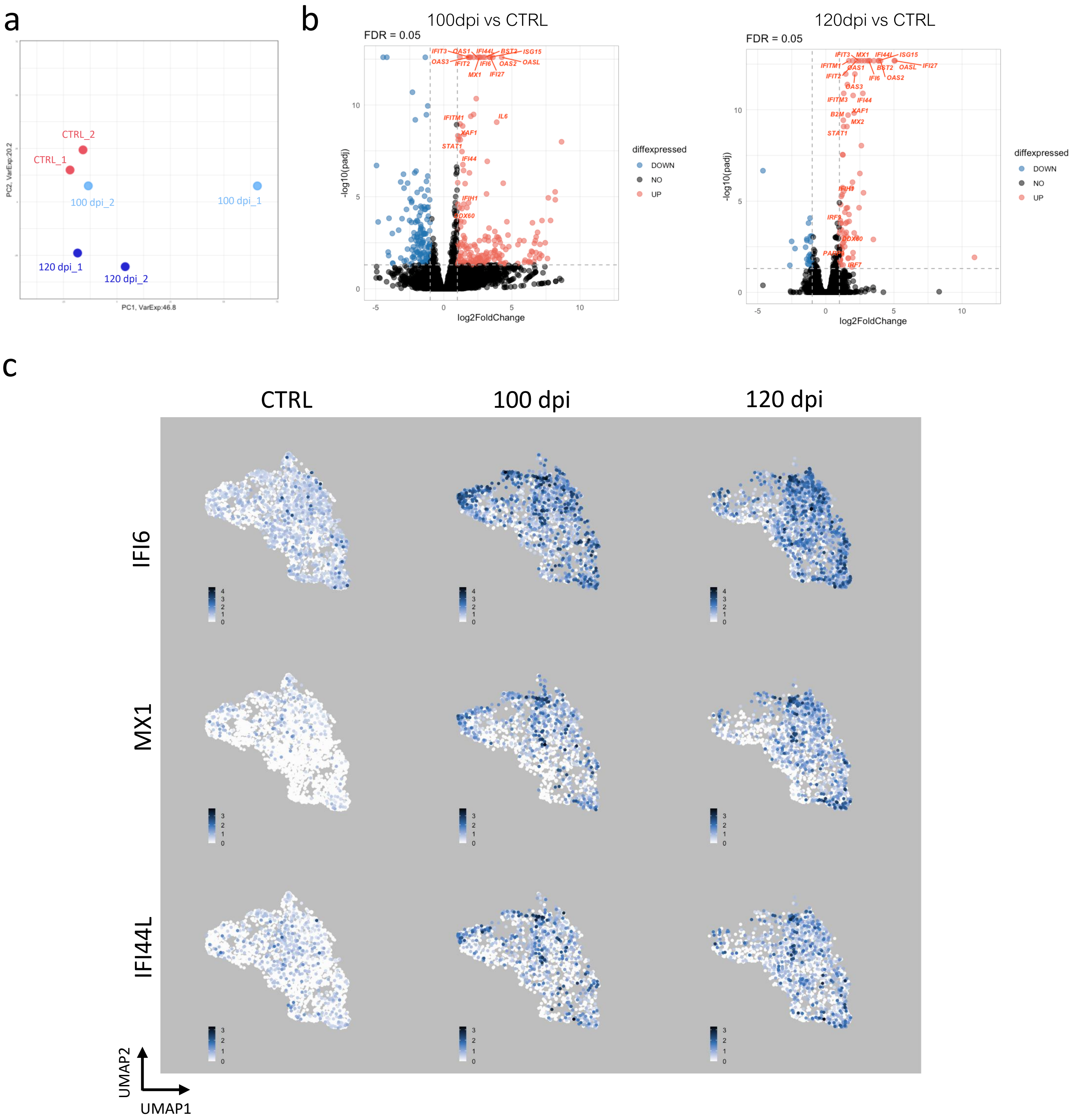
a
b
c
CTRL_2
CTRL_1
100 dpi_2
100 dpi_1
120 dpi_1
120 dpi_2
PC2, VarExp:20.2
PC1, VarExp:46.8
100dpi vs CTRL
FDR = 0.05
120dpi vs CTRL
FDR = 0.05
-log10(padj)
log2FoldChange
diffexpressed
DOWN
NO
UP
CTRL
100 dpi
120 dpi
IFI6
MX1
IFI44L
UMAP2
UMAP1

Figure_S12

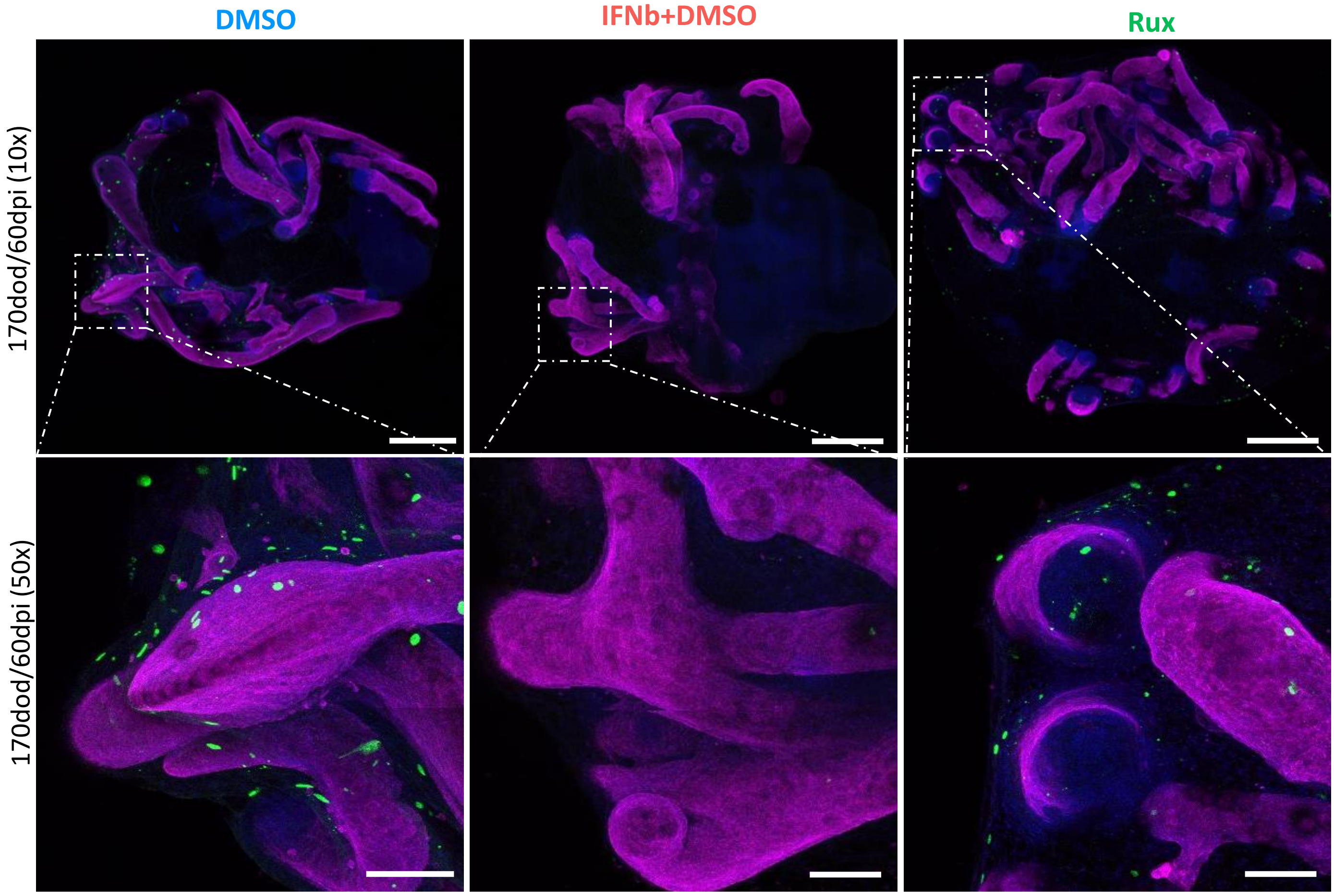
