## Supplementary Table for "Merkel cell polyomavirus infection and persistence modelled in skin organoids"

**Table 1 | Antibody list**

| Primary antibodies | Markers for: | clone n. | Cat. No. | source | species | Isotype | IHC/IF FFPE | IF on cells | WM on SkOs |
|---|---|---|---|---|---|---|---|---|---|
| E-Cadherin | Epithelium | 36/E-Cadherin | 610181 | BD Biosciences | Mouse | IgG2a | 1:100 | | 1:50 |
| Ki67 | Proliferating cells | B56 | 550609 | BD Biosciences | Mouse | IgG1 | 1:100 | | 1:100 |
| KRT15 | | LHK15 | sc-47697 | Santa Cruz | Mouse | IgG2a | | | 1:50 |
| KRT17 | Periderm, epidermis & outer root sheath | D-4 | sc-393091 | Santa Cruz | Mouse | IgG1 | | | 1:100 |
| KRT20 | Merkel cells | D9Z1Z | 13063S | Cell Signaling | Rabbit | IgG | | | 1:100 |
| LHX2 | Hair placodes & bulge | | ABE1402 | Millipore | Rabbit | IgG | | | 1:750 |
| LT | MCPyV LT | Ab3 | | James DeCaprio, DF/HCC | Mouse | IgG1 | 1:250 | | 1:200 |
| LT | MCPyV LT | Cm2B4 | sc-136172 | Santa Cruz | Mouse | IgG2b | | 1:500 | |
| PDGFRα | Dermal fibroblasts | D13C6 | 5241 | Cell Signaling | Rabbit | IgG | 1:100 | | 1:50 |
| SOX2 | Dermal papilla/condensates, Merkel cells, Melanocytes | O30-678 | 561469 | BD Biosciences | Mouse | IgG1 | | | 1:100 |
| TUBB3 | Neurons | TUJ1 | 801202 | BioLegend | Mouse | IgG2a | | | 1:100 |
| VP1 | MCPyV VP1, capsid protein | | | Christopher B. Buck, NCI | Rabbit | IgG | | 1:1000 | 1:1000 |

| Secondary ab target | Isotype | species reactivity | Cat.No. | source | fluorophore | IHC/IF FFPE | IF on cells | WM on SkOs |
|---|---|---|---|---|---|---|---|---|
| Anti-mouse | IgG | Goat | A-11004 | Thermo Fisher Scienfitic | A-568 | | | 1:2000 |
| Anti-mouse | IgG | Goat | A-11001 | Thermo Fisher Scienfitic | A-488 | 1:600 | | 1:2000 |
| Anti-mouse | IgG1 | Goat | A-21240 | Thermo Fisher Scienfitic | A-647 | | | 1:2000 |
| Anti-mouse | IgG1 | Goat | A-21121 | Thermo Fisher Scienfitic | A-488 | | 1:600 | 1:2000 |
| Anti-mouse | IgG2a | Goat | A-21134 | Thermo Fisher Scienfitic | A-568 | | 1:600 | 1:2000 |
| Anti-rabbit | IgG | Goat | A-11008 | Thermo Fisher Scienfitic | A-488 | | | 1:2000 |
| Anti-rabbit | IgG | Goat | A-11036 | Thermo Fisher Scienfitic | A-568 | 1:600 | | 1:2000 |
| Anti-rabbit | IgG | Goat | A-21244 | Thermo Fisher Scienfitic | A-647 | | 1:600 | 1:2000 |
| Hoechst | | | H3570 | Thermo Fisher Scienfitic | | | | 1:1000 |
